# Complementary remodeling strategies distinguish human subcutaneous and omental adipose tissue

**DOI:** 10.64898/2026.07.18.739362

**Authors:** Gerelt-Od Khenmedekh, Dae Hoon Kim, Seung-Myoung Son, Young Chul Kim, Myoung Won Son, Jong Hyuk Yun

## Abstract

**Background:** Subcutaneous adipose tissue (SAT) and visceral adipose tissue (VAT) differ in their metabolic risk, but whether they retain distinct transcriptional identities and remodeling programs in adult humans remains unclear.

**Methods:** Bulk RNA sequencing was performed on 29 adipose specimens from 19 patients, including 10 paired SAT–VAT samples, along with baseline CT-derived depot area and attenuation measurements. The findings were compared with Masson trichrome staining and CD68 histology in an independent cohort of 30 patients and validated using GTEx adipose tissue data and an external human single-nucleus atlas.

**Results:** SAT and VAT showed distinct transcriptomic identities. SAT was enriched for a mesenchymal patterning program characterized by **TBX15** and **SHOX2**, whereas omental VAT exhibited a mesothelial–stromal signature marked by **UPK3B**. These depot-specific identity signals remained significant after adjusting for BMI, age, and measured cellular signatures. SAT area correlated with extracellular matrix remodeling, whereas VAT area correlated with vascular-hypoxia signaling. These associations were attenuated after BMI adjustment, indicating that remodeling was linked to overall adiposity. In an independent histological cohort, SAT exhibited substantially greater fractional fibrosis than VAT, whereas VAT demonstrated a markedly higher storage-to-scaffold index. Depot-associated transcriptional effects were independently reproduced in the external datasets.

**Conclusions:** Human SAT and omental VAT retain distinct tissue identities and exhibit complementary remodeling strategies. SAT preferentially adopts a mesenchymal–ECM scaffold program, whereas VAT favors mesothelial–stromal and vascular remodeling programs. These findings support a storage-versus-scaffold framework for adaptation of human adipose tissue to chronic excess energy.

## 1. Introduction

White adipose tissue is organized into anatomically and functionally distinct depots. Subcutaneous (SAT) and visceral fat (VAT) differ in their association with insulin resistance and cardiometabolic risk, with visceral accumulation generally carrying greater metabolic liability [1]. A long-standing question is whether these depots differ merely because of their anatomical location and local microenvironment, or because they constitute intrinsically distinct tissue programs that respond differently to energy surplus [1, 2].

Human and rodent studies have reported depot-biased expression of developmental and positional genes, including HOX cluster members TBX15, SHOX2, and EN1, and have proposed that positional identity is partially maintained in adulthood [3–6]. TBX15, a mesodermal T-box transcription factor, is consistently higher in subcutaneous fat than in visceral fat and correlates with adiposity and fat distribution [1, 3, 7]. In parallel, omental VAT is covered by a mesothelial lining, and markers such as UPK3B, KRT19, MSLN, and WT1 identify mesothelial and mesothelial-associated stromal populations rather than adipocytes per se [2, 8, 9]; KRT19 marks omental mesothelium and is essentially absent from subcutaneous fat [8].

Several caveats apply to bulk tissue interpretation. First, signatures built from a few handpicked genes (e.g., selected HOX loci) may not represent the broader transcriptional program. Second, because VAT specimens include a mesothelial surface and differ from SAT in stromal-vascular composition, bulk depot differences may partly reflect cell-type proportions rather than adipocyte-intrinsic programs [8, 10, 11]. Third, transcriptional “identity” is not equivalent to a distinct embryonic lineage.

In this study, we adopt a deliberately conservative approach using paired human specimens. We ask whether (i) depot identity is robust to gene selection and can be adjusted for measured cellular signatures and BMI, and (ii) whether depot identity is accompanied by distinct tissue-level remodeling programs that are evident in histology and CT. We show that the most robust identity signal is mesenchymal (TBX15/SHOX2) for SAT and mesothelial–stromal for omental VAT, rather than a broad HOX program, and that the two depots exhibit divergent ECM-stiffening versus vascular–stromal remodeling in association with adiposity. We frame these findings as evidence of depot-associated transcriptional identity and remodeling, while deferring questions regarding lineage, epigenetics, and postsurgical outcomes for future studies.

## 2. Methods

### 2.1 Cohort and specimens

Paired subcutaneous and omental adipose tissue specimens were obtained from patients undergoing abdominal surgery at Chungbuk National University Hospital. This study was approved by the Institutional Review Board of Chungbuk National University Hospital (IRB No. CBNU-2025-A-0174; expedited review; approved 2025-12-29) and conducted in accordance with the Declaration of Helsinki. A waiver of informed consent was granted for the retrospective analysis of archived specimens. Specimens were collected intraoperatively, snap-frozen or fixed, as appropriate, and stored until analysis. Because the surgical population primarily comprised patients with gastrointestinal cancer and undergoing metabolic surgery, the recruited cohort was enriched for individuals at the lean and obese ends of the BMI spectrum, with relatively few overweight (BMI 23–29 kg/m²) individuals. This distribution is reflected in the observed BMI distribution (Section 3.4).

### 2.2 Depot assignment

The authoritative depot label was based on operative/pathology tissue annotation (subQ = SAT, omentum = VAT). The identity genes used as readouts (TBX15, SHOX2, and UPK3B) were not used to assign labels, thereby avoiding circularity. Sample-flow accounting and low-confidence sensitivity analyses are provided in the Supplementary Methods and Supplementary Table S4.

### 2.3 RNA sequencing and processing

Total RNA was sequenced by a commercial provider (Macrogen, Seoul, Republic of Korea) using a capture-based protocol (Agilent SureSelectXT RNA Direct) suitable for partially degraded RNA, which is typical of adipose tissue (paired-end, 151 bp; 29 samples; 16,493 genes retained). Depot differential expression analysis used edgeR (with TMM normalization; using exactTest at |fold change| ≥ 2 and unadjusted p < 0.05 as an exploratory discovery threshold); a complementary within-patient paired comparison (10 paired samples) is reported in Supplementary Table S2, and the external-validation discovery set was defined by an FDR < 0.05 and |log2FC| > 1 (Section 2.9). Full RNA extraction, QC, and processing details are provided in the Supplementary Methods.

### 2.4 Module scores

Gene modules were defined a *priori* (identity: mesenchymal patterning and mesothelium; remodeling: ECM stiffening, vascular-hypoxia, and macrophages). Each module score was calculated as the mean across the member genes of the cohort-wide per-gene z-scores. The analyzed module compositions are listed in Supplementary Table S1 and the Supplementary Methods.

### 2.5 Statistical analysis

Paired SAT–VAT comparisons were performed using the exact Wilcoxon signed-rank test on within-patient differences with rank-biserial effect sizes reported. Group comparisons were performed using the Mann–Whitney U test. Linear mixed-effects models (including a patient random intercept) were used to test the depot effect with adjustment for measured cell composition scores and separately for BMI and age (coefficients are provided in Supplementary Table S3). Principal component analysis, a leave-one-patient-out classifier, and CT–RNA partial correlation analysis were performed as described in the Supplementary Methods. Analyses were performed in Python (version 3.11.7; Python Software Foundation, Beaverton, OR, USA) using a reproduction package that regenerates every reported statistic (see Data Availability).

### 2.6 Primary integrative analyses

Two directional hypotheses were designated primary: SAT area ↔ SAT ECM module and VAT area ↔ VAT vascular–hypoxia module; the independent histologic cohort provided group-level (rather than within-patient) support, as only one patient overlapped between cohorts. All other genes, including those associated with CT and metabolism, were also examined.

Two 2-D CT–histology surrogates were defined per depot from the single-slice CT area (A, cm²) and histologic fibrosis area fraction (f, %): the area-weighted fibrosis index = A × (f/100), and the storage-to-scaffold index = A/f (cm² per percentage point of fibrosis; higher values denote more adipose area per unit fibrosis). These surrogates were computed from a single CT slice and a separate histological section and do not represent direct measurements of tissue volume, storage capacity, or collagen mass. Accordingly, they are referred to as indices throughout the study (Supplementary Methods).

### 2.7 Histology (independent cohort)

Histological analysis was performed on an independent cohort (n = 30 paired samples; overlap with the RNA cohort = one patient). Masson trichrome staining and CD68 immunohistochemistry were quantified using QuPath v0.5.1 with a supervised fibrosis classifier trained on a representative subcutaneous section and applied uniformly across all SAT and VAT specimens to quantify the fibrosis area fraction (%) (scale bars, 300 μm).

Because a single-depot-trained classifier may be conservative for VAT, the depot fibrosis comparison is considered directionally robust in direction. Staining, imaging, and classifier details and study limitations are provided in the Supplementary Methods.

### 2.8 CT analysis (baseline)

Baseline non-contrast abdominal CT, obtained as part of routine preoperative care within 30 days before surgery, was segmented at the L3 level using 3D Slicer (v5.10.0; −190 to −30 HU) to obtain SAT and VAT cross-sectional areas (cm²) and mean/median attenuation (HU). Full segmentation details are in Supplementary Methods.

In the histological cohort (n = 30), the CT measurements of area and HU were complete and within the expected adipose tissue range; a few RNA cohort specimens showed atypically high HU (as flagged in Supplementary Table S1b). HU-based analyses should therefore be interpreted as exploratory. Postoperative and metabolic outcomes were not analyzed.

### 2.9 External validation (public datasets)

To assess generalizability, the depot programs were compared with two external human datasets: GTEx v8 adipose–subcutaneous tissue (n = 663) and adipose–visceral/omental tissue (n = 541) [12], and the Emont et al. single-nucleus atlas [11] (137,684 nuclei; aggregated to donor × depot × cell-type pseudobulk). Cross-dataset concordance was quantified using the Spearman correlation of depot log2 fold-changes across shared genes and the proportion of discovery DEGs with a matching direction. The full pipeline is in Supplementary Methods. Analysis scripts and processed matrices are provided (see Data availability).

## 3. Results

### 3.1 Global transcriptomic separation of SAT and omental VAT

SAT and VAT specimens (n = 11 and 18, respectively) were separated along the first principal component of the most variable genes (rank-biserial correlation 0.83, p = 0.0002). This separation was maintained after excluding mesothelial, immune, endothelial, and erythroid genes (rank-biserial correlation 0.80, p = 0.0004), indicating that global depot separation is not solely driven by these cell type-associated transcripts (Figure 1). Paired differential expression analysis identified depot-biased genes, consistent with previous reports, including SAT-enriched positional/mesenchymal and VAT-enriched mesothelial/immune genes (Supplementary Table S2).

**Figure 1.**
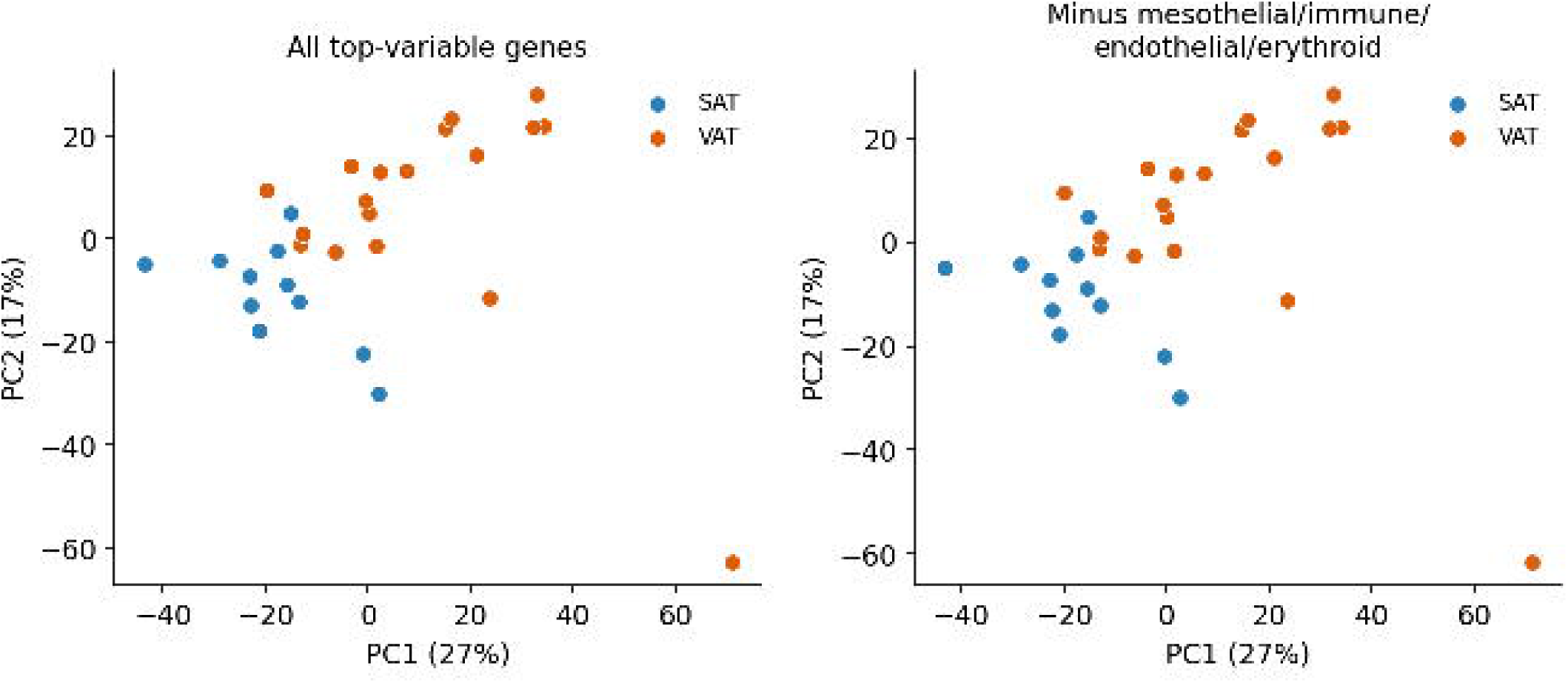
Cohort and global depot separation. (A) Study design and paired SAT–VAT sampling (10 pairs among 29 specimens; SAT n = 11, VAT n = 18). (B) PCA on the top 2,000 variable genes showing SAT–VAT separation (rank-biserial 0.83, p = 0.0002). (C) PCA after excluding mesothelial/immune/endothelial/erythroid genes, with separation retained (rank-biserial 0.80, p = 0.0004). Points are individual samples; the single VAT outlier is retained (sensitivity analysis in Supplementary Figure S7).

### 3.2 The robust identity axis is mesenchymal (SAT) and mesothelial–stromal (VAT), not broad HOX

Among the identity modules, the mesenchymal-patterning signature was higher in SAT and remained significant at the single-gene level; TBX15 and SHOX2 each showed paired SAT– VAT differences with p = 0.002 (Figure 2). Conversely, the mesothelial signature was higher in omental VAT and robust to single-gene testing (UPK3B paired, p = 0.002). These directions are concordant with prior human studies demonstrating subcutaneous-dominant *TBX15/SHOX2* [3–6] and omental-restricted *KRT19/UPK3B* mesothelial markers [8, 9].

**Figure 2.**
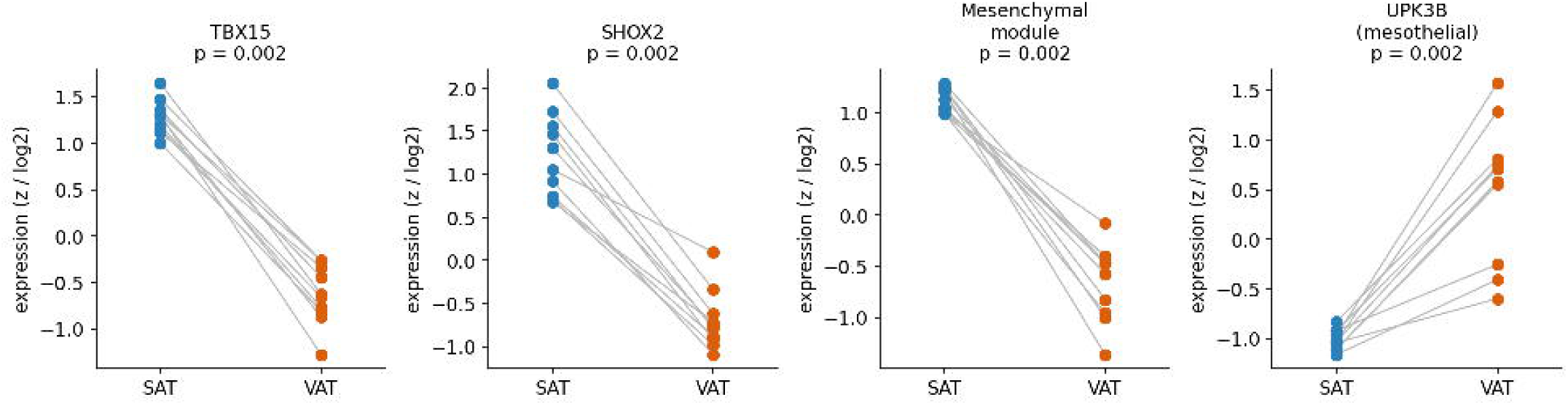
Robust depot-associated identity. Within-patient paired plots (10 pairs) for TBX15 and SHOX2 (SAT-high), the mesenchymal-patterning module, and UPK3B (VAT-high, mesothelial); each point is a sample, lines connect paired specimens, and p is paired Wilcoxon. Values are cohort-wide per-gene z-scores (single genes) or module z-scores. TBX15/SHOX2/UPK3B were not used for depot assignment (Methods 2.2). HOX analyses are in Supplementary Figure S1.

In contrast, a broad HOX program did not robustly distinguish the depots; significance depended on single genes and disappeared when all detected HOX genes were included, with the HOXA cluster even reversing direction (Supplementary Figure S1; Supplementary Results). Therefore, we interpreted the most robust identity signal as being carried by the mesenchymal-patterning genes (*TBX15/SHOX2*) and the VAT mesothelial–stromal program, and not by a broad HOX axis.

### 3.3 The SAT mesenchymal signal persists after adjustment for measured cellular signatures and BMI

The SAT mesenchymal signal was largely retained after adjusting for mesothelial and macrophage signature scores (approximately 92% of the coefficient; p < 0.0001) and remained highly significant after adjusting for BMI and age (p < 0.0001) (Figure S4), indicating that depot-associated identity is not fully explained by the measured cell composition or by adiposity. We interpret this conservatively and do not claim independence from unmeasured stromal, vascular, or progenitor populations (Supplementary Results).

### 3.4 Depot-associated remodeling programs are associated with adiposity

SAT cross-sectional area correlated with the ECM cross-linking/stiffening module (ρ = 0.77, p = 0.005), whereas VAT area correlated with the vascular–hypoxia module (ρ = 0.57, p = 0.013) but not with macrophage module content (ρ = 0.03, p = 0.91) (Figure S6). These associations were attenuated after BMI adjustment, consistent with depot remodeling tracking overall adiposity rather than reflecting a unique local effect (Supplementary Results; Supplementary Table S3). The biologically important observation is that the two depots engage distinct remodeling modules as adiposity increases (Figure 4). Within SAT, *LOX* expression was higher in the obese subgroup than in the lean subgroup (reflecting the bimodal BMI distribution; Supplementary Figure S2).

### 3.5 Histologic and imaging correlates

In an independent histologic cohort (n = 30; overlap with the RNA cohort = 1 patient), SAT showed markedly higher interstitial fibrosis than VAT (17.8% vs 4.2%, paired Wilcoxon p ≈ 3×10⁻⁶) (Figure 3), providing independent, tissue-level support for the SAT ECM/stiffening program. The CD68-positive area fraction was very low in both depots, and given its small magnitude and the absence of any VAT area and the macrophage RNA module, it is not interpreted as histological support for a VAT macrophage axis (Supplementary Results; Supplementary Figure S5). Because the histological cohort was independent of the RNA cohort (overlap = 1 patient), these observations should be interpreted as orthogonal biological support rather than patient-level validation.

**Figure 3.**
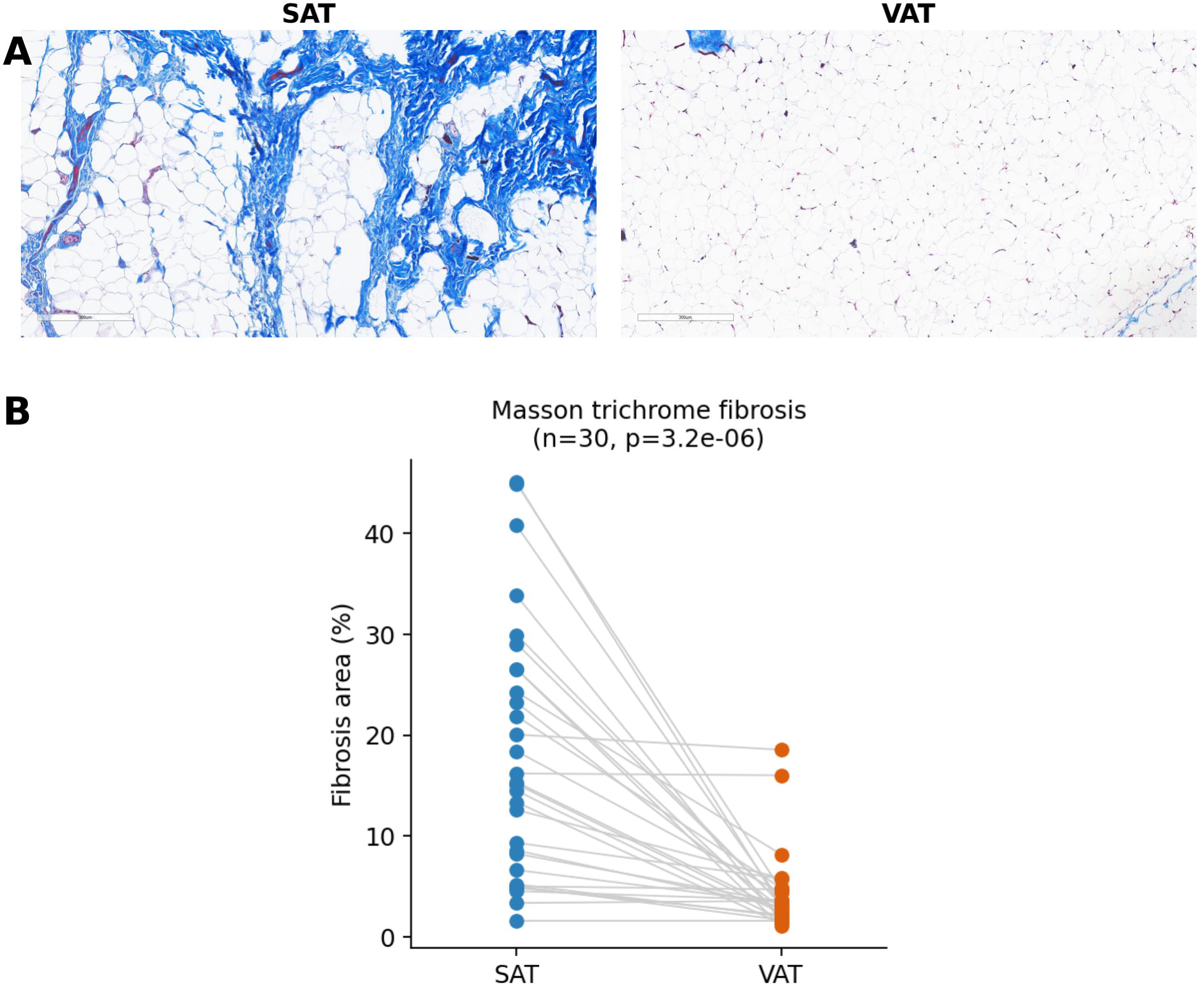
Histologic fibrosis in an independent cohort. (A) Representative Masson trichrome sections for SAT and VAT (100× total magnification, 10× objective; scale bar 300 μm; representative of median fibrosis, Methods 2.7). SAT shows more abundant collagen deposition (blue) than VAT. (B) Paired quantification (n = 30): SAT fibrosis markedly exceeds VAT (17.8% vs 4.2%, paired Wilcoxon p ≈ 3×10⁻⁶). CD68 immunohistochemistry, which showed only a small absolute depot difference, is presented in Supplementary Figure S5.

**Figure 4.**
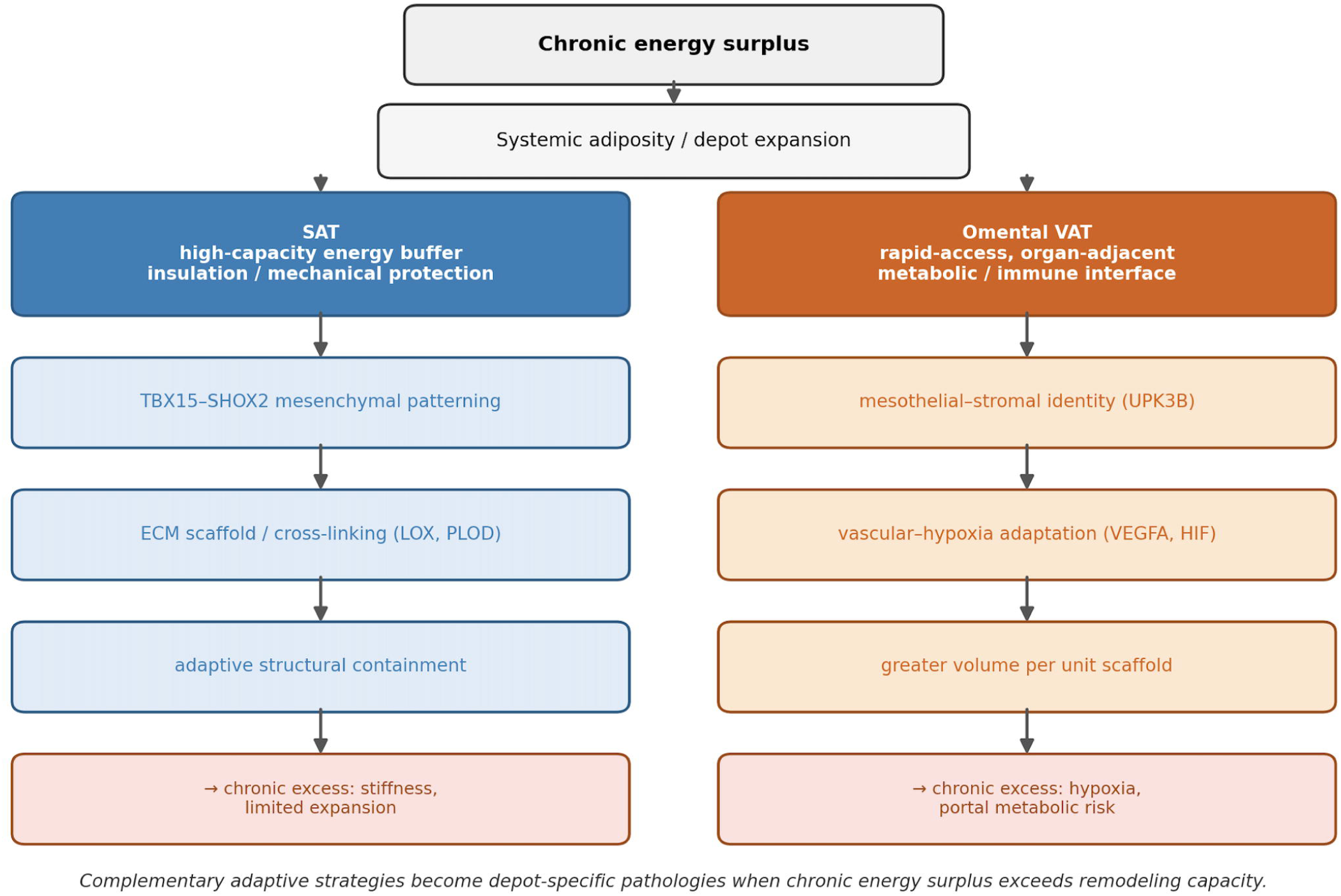
Integrated model. On a background of fixed SAT mesenchymal versus VAT mesothelial–stromal identity, increasing adiposity is associated with depot-specific remodeling: SAT engages ECM containment/fibrosis (histologically supported) as a high-capacity but scaffold-constrained storage buffer, whereas VAT engages an area-associated vascular–hypoxia program (not histologically confirmed) with a higher storage-to-scaffold index (greater adipose area per unit fractional fibrosis). Complementary adaptive strategies may become depot-specific pathologies when chronic energy surplus exceeds remodeling capacity.

### 3.6 SAT and VAT differ in the relationship between scaffold and storage

Because the histologic cohort had completely paired CT depot areas for all 30 patients, we directly related fibrosis density to depot size (Figure 5). Fibrosis fraction was inversely related to depot area in SAT (ρ = −0.45, p = 0.012), and high-fibrosis SAT specimens had significantly smaller SAT area than low-fibrosis specimens (median 111 vs 162 cm², p = 0.003), consistent with a scaffold-associated ceiling on subcutaneous expansion. Despite this higher fractional fibrosis in SAT, omental VAT showed a several-fold more adipose area per unit of fractional fibrosis (storage-to-scaffold index 46.5 vs 8.7 cm² per %; VAT > SAT in 28/30 patients; paired p < 0.0001). Robustness analyses (outlier removal; male and non-diabetic subgroups), weak SAT-fibrosis–to–VAT/SAT-ratio tendencies that did not survive full covariate adjustment, and CT attenuation (HU) as a depot-associated imaging phenotype distinct from fibrosis are reported in the Supplementary Results (Supplementary Figure S8).

**Figure 5.**
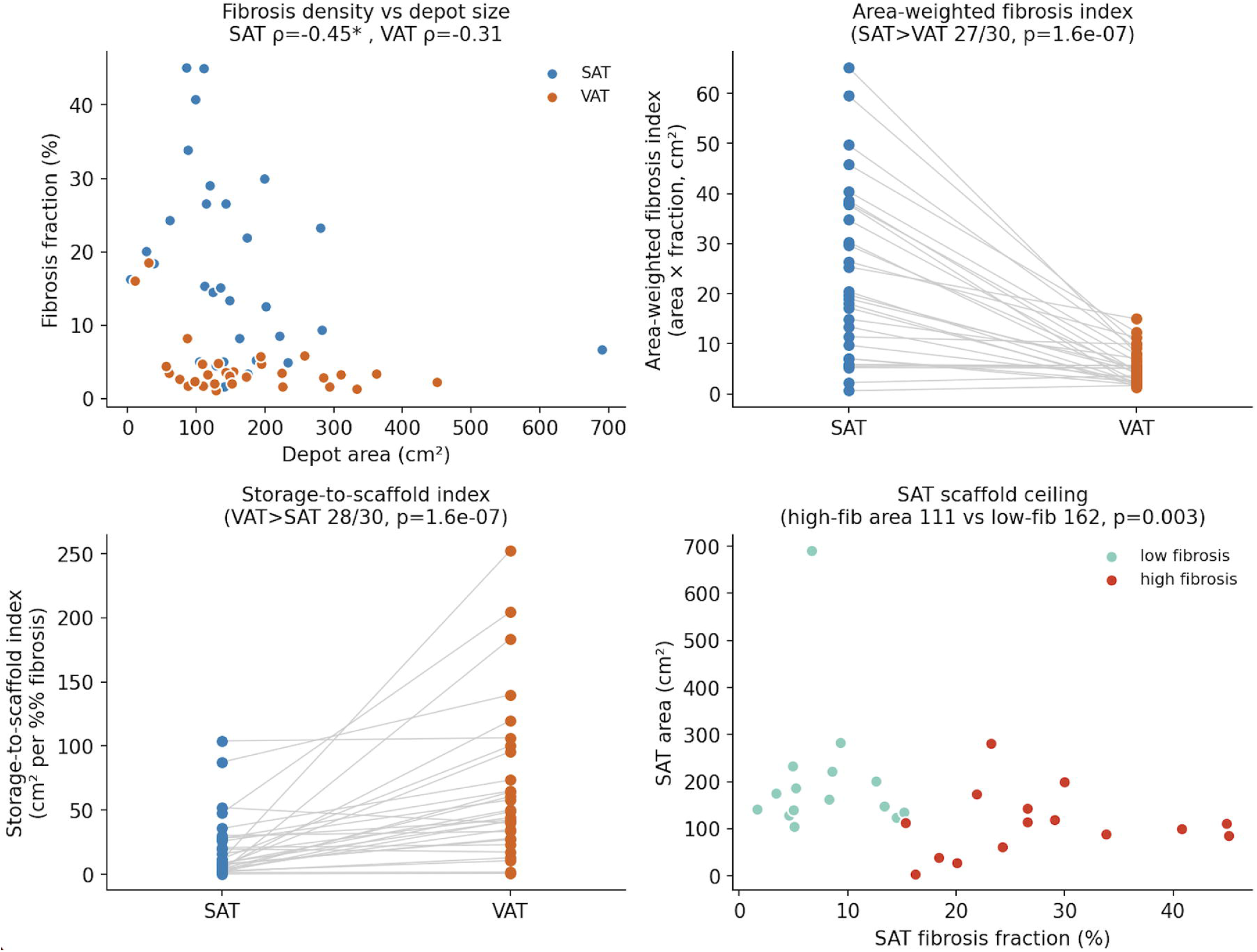
Storage-versus-scaffold relationship differs between depots (independent cohort, n = 30). (A) Fibrosis fraction versus depot area: inverse in SAT (ρ = −0.45, p = 0.012; robust to outlier trimming and present in male and non-diabetic subgroups) and weakly inverse in VAT. (B) Area-weighted fibrosis index (area × fraction, cm²) is higher in SAT (SAT > VAT in 27/30; p < 0.0001). (C) Storage-to-scaffold index (area per percentage-point of fibrosis, cm² per %) is far higher in VAT (median 46.5 vs 8.7; VAT > SAT in 28/30; p < 0.0001; robust to removing the most extreme value, 27/29). (D) SAT scaffold ceiling: high-fibrosis SAT specimens have smaller SAT area than low-fibrosis specimens (median 111 vs 162 cm², p = 0.003). The SAT fibrosis–VAT/SAT ratio association (storage diversion) was borderline and did not survive full covariate adjustment; it is treated as hypothesis-generating (Section 3.6, Supplementary Table S5).

### 3.7 Depot transcriptional architecture is reproducible in external datasets

To test whether depot separation reflected reproducible biology rather than cohort-specific noise, we compared genome-wide depot effect sizes across two independent datasets (Figure 6; Table S6). Effect sizes were concordant with GTEx adipose tissues (Spearman ρ = 0.325 across 13,958 genes; 360/403 strong discovery DEGs showed directional concordance, 89.3%) and, more strongly, with donor-level pseudobulk from the Emont single-nucleus atlas [11] (ρ = 0.511; 362/374 genes were directionally concordant, 96.8%). In addition, 341 genes were significant in both our cohort and GTEx with concordant direction. Replicated genes recapitulated both identity axes—SAT developmental/positional (*TBX15, IRX2, and SHOX2*) and VAT mesothelial/epithelial (*MUC16, UPK3B, and KRT19*)—establishing that depot separation is generalizable to independent cohorts.

**Figure 6.**
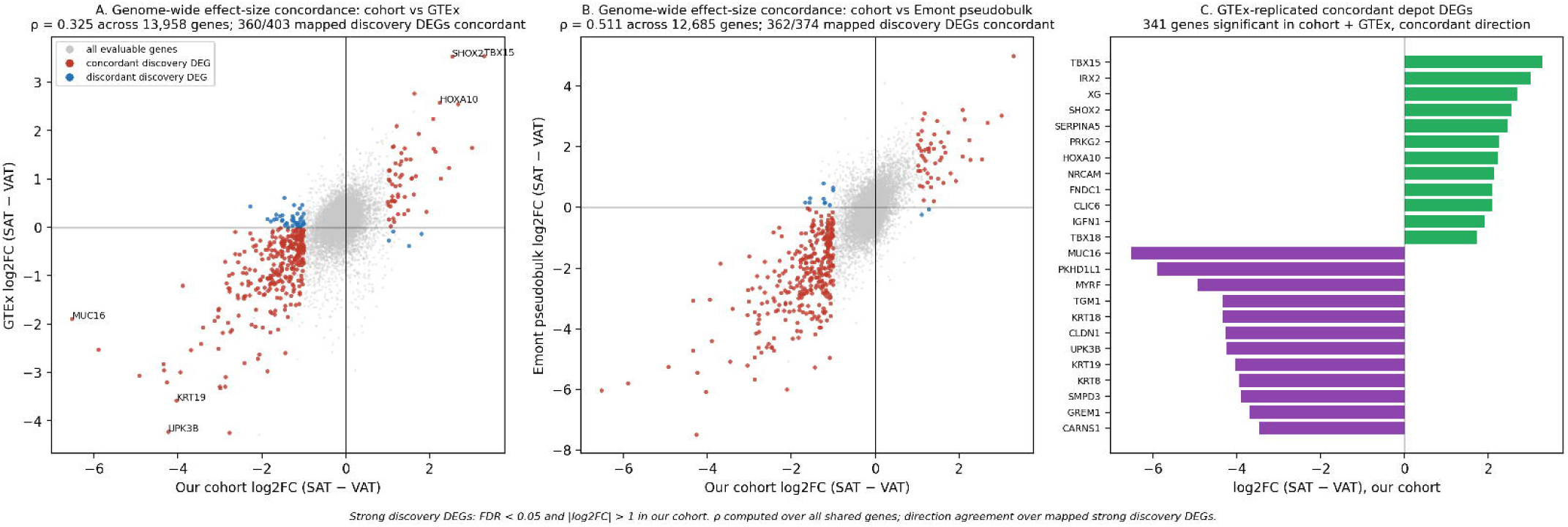
Genome-wide depot effects are reproducible in external datasets. (A) Depot log2 fold-changes (SAT − VAT) in our cohort versus GTEx adipose tissues (subcutaneous n = 663, omental n = 541); gray, all evaluable genes; red, strong discovery DEGs with concordant direction; blue, strong discovery DEGs with discordant direction (strong discovery DEGs: FDR < 0.05 and |log2FC| > 1 in our cohort). Spearman ρ = 0.325 across 13,958 shared genes; 360/403 mapped discovery DEGs concordant. (B) The same comparison against donor-level pseudobulk from the Emont single-nucleus atlas [12]; ρ = 0.511 across 12,685 shared genes; 362/374 mapped discovery DEGs concordant. (C) GTEx-replicated concordant depot DEGs (341 genes significant in our cohort and GTEx with concordant direction): top SAT-enriched (green) and VAT-enriched (purple) genes by effect size. Full-resolution figure files are provided separately (see Reproduction package).

At single-nucleus resolution, fibrillar collagen expression (COL1A1, COL3A1, and COL6A3) was concentrated in adipose stem/progenitor cells (ASPCs), whereas LOX was distributed across stromal and vascular cell types (Figure S9), indicating that the SAT ECM signal is a progenitor– and stroma-associated program rather than an adipocyte-intrinsic one. Thus, the public datasets validate the depot architecture, whereas our cohort uniquely connects that architecture to BMI, CT adiposity, and histological fibrosis (Supplementary Results).

## 4. Discussion

In paired human specimens, subcutaneous and omental adipose tissues maintained distinct transcriptional identities and engaged in divergent remodeling programs. The most robust identity signal was mesenchymal (TBX15/SHOX2) for SAT and mesothelial–stromal for omental VAT; however, a broad HOX program was not supported. The depot-associated identity remained significant after adjusting for BMI, age, and measured cellular signatures, whereas the depot-specific remodeling associations tracked adiposity, and SAT showed higher fibrosis than VAT in an independent histological cohort. These depot effect sizes were reproducible in GTEx and in an independent single-nucleus atlas (Spearman ρ = 0.33 and 0.51; 341 GTEx-replicated concordant genes), and single-nucleus data localized the SAT ECM signal to stromal/progenitor rather than adipocyte compartments. This addresses the principal limitation of a small cohort while leaving the link to BMI, CT adiposity, and histologic fibrosis as the distinct contribution of this study.

### Mechanistic interpretation

The SAT-dominant mesenchymal patterning signature (TBX15 and SHOX2) is consistent with a body of work establishing mesodermal factors as stable, depot-biased markers of subcutaneous fat that influence preadipocyte behavior and adipogenic capacity [1, 3–7, 13]. TBX15 marks a glycolytic pre-adipocyte/adipocyte subpopulation and is inversely related to adiposity [7, 13], providing a plausible link between fixed positional identity and depot-specific expandability. The coupling we observed between SAT identity and an ECM cross-linking/stiffening program (LOX, POSTN, PLOD, P4HA) fits mechanistic models in which lysyl oxidase-mediated collagen cross– linking, rather than bulk collagen synthesis, governs matrix stiffness and constrains healthy expansion [14–17]. LOX-driven cross-linking increases ECM rigidity and is associated with insulin resistance, while LOX inhibition reduces matrix stiffness and improves metabolic parameters in obese models [17]. On the visceral side, the omental mesothelial–stromal signature (UPK3B, KRT19, TCF21) reflects the mesothelial lining and mesothelial-associated stromal characteristics of the omentum rather than a distinct visceral adipocyte identity [2, 8, 9] Mesothelial cells display plasticity and can undergo a mesothelial-to-mesenchymal transition, thereby contributing to the stromal milieu and omental responses to inflammation and cancer [2, 9]. Accordingly, we described the VAT axis as an area-associated vascular-hypoxia transcriptional program [18–20] rather than a histologically confirmed phenotype.

### A storage-versus-scaffold framework

These molecular and histological differences are consistent with the physiological division of labor between the two depots. Subcutaneous fat is not a passive reservoir but a protective energy buffer that sequesters surplus lipids in a high-density form and shields the liver, muscle, and pancreas from ectopic lipid exposure; severe insulin resistance and hepatic steatosis observed in lipodystrophy illustrate what happens when this buffering capacity is lost [1, 21]. As the largest and most expandable subcutaneous compartment, SAT is well suited for long-term, high-capacity storage; its expansion requires a proportionate extracellular scaffold to support enlarging lobules, distribute tissue tension, and prevent adipocyte rupture and necrosis. Therefore, SAT fibrosis need not be interpreted as purely pathological scarring; it may begin as an adaptive scaffold and become maladaptive, imposing stiffness and restricting expandability only when it exceeds a threshold [14, 15, 22, 23]. Our histological data directly support this interpretation; fractional fibrosis was inversely related to the SAT area, and high-fibrosis SAT specimens were significantly smaller, consistent with a scaffold-associated ceiling on subcutaneous expansion. Omental VAT, by contrast, showed several-fold more adipose area per unit of fractional fibrosis (storage-to-scaffold index) and engages in vascular-hypoxia rather than a ECM-stiffening program, consistent with a more dynamic, organ-adjacent depot with high vascular-stromal plasticity, rapid lipolytic responsiveness, and direct portal connection to the liver [1, 2, 18]. We observed a weak, direction-consistent tendency for higher SAT fibrosis to accompany a higher VAT/SAT ratio, but this did not survive full covariate adjustment. Therefore, whether a scaffold-constrained subcutaneous compartment actively diverts storage toward the visceral depot remains a hypothesis for prospective testing rather than a demonstrated mechanism.

### Clinical and physiological implications

The finding of higher fractional fibrosis in SAT than in VAT was superficially unexpected, as visceral fibrosis is often emphasized in metabolic diseases [22]. However, subcutaneous fibrosis is itself associated with insulin resistance and limited fat-mass plasticity [14, 15, 22], and the depot-specific coupling of SAT identity to an ECM-stiffening program supports the view that SAT and VAT deploy fundamentally different structural strategies rather than indicating that one depot is simply “healthier.” The omental vascular–hypoxia program aligns with the higher metabolic liability historically attributed to visceral fat and its portal delivery of free fatty acids and inflammatory signals [1, 18]. Accordingly, reframing obesity therapy, accordingly, the goal may be less about indiscriminate loss of fat mass than about restorating of healthy depot-specific remodeling, preserving safe subcutaneous storage while relieving pathological ECM stiffening and reducing visceral hypoxic–vascular dysfunction. Depot-specific interventions are conceivable on this basis: antifibrotic or LOX-targeted strategies to normalize SAT ECM turnover and restore expandability [17], versus approaches aimed at VAT perfusion, hypoxia, and stromal inflammation. Because bariatric surgery reduces LOX expression and collagen cross-linking and is accompanied by ECM remodeling and neovascularization associated with metabolic improvement [24], baseline depot-specific remodeling states—captured by SAT ECM/fibrosis and VAT vascular-hypoxia measures— could, in principle, predict post-surgical weight loss and metabolic response. This is an explicit motivation for the longitudinal study we are undertaking. CT attenuation may add a complementary imaging phenotype (depot HU tracked depot size and differed between SAT and VAT after BMI adjustment in this cohort) that, if its biological basis and sensitivity to segmentation are validated in larger cohorts, could serve as a non-invasive readout distinct from depot volume.

**From an evolutionary perspective, although our cross-sectional data motivate but cannot prove—regional adipose specialization may reflect complementary survival functions: subcutaneous fat for high-capacity, protected long-term storage and visceral fat for rapid substrate mobilization and local immune–vascular responses. Under chronic energy surplus these initially adaptive strategies may be overrun, converting SAT scaffold formation into restrictive fibrosis and VAT vascular adaptation into hypoxic, metabolically hazardous remodeling**.

### Limitations

The sample was modest (11 SAT, 18 VAT; 10 pairs), so the results were effect size– and direction-oriented and hypothesis-generating. Bulk RNA cannot resolve whether identity signals are adipocyte-intrinsic. Markers such as UPK3B reflect a mesothelialstromal niche rather than visceral adipocyte identity, and we framed VAT identity accordingly [8, 10]. The bimodal BMI distribution precludes reliable dose– response inference for remodeling markers, and CT–RNA associations are confounded by adiposity. We studied omental VAT only; the mesenteric and epiploic visceral depots may have differed. The cohort was Korean and had a single center; therefore, ethnic and site generalizability requires replication. Input RNA was partially degraded (DV200-based QC) and libraries were prepared with a capture-based protocol (SureSelect RNA Direct) suited to such material; depot contrasts should be robust to this, and absolute expression levels and low-abundance transcripts should be interpreted with corresponding caution. Histology is derived from an independent cohort, providing group-level rather than within-patient support, and RNAhistology cohorts overlap by a single patient. The fibrosis classifier was trained on a single subcutaneous slide and applied uniformly; while this supports the direction of the SAT > VAT difference, the absolute VAT fibrosis fraction may be conservative, and a both-depot-trained classifier with pathologist concordance is planned. Finally, the design was cross-sectional, and causality could not be established. External comparison with GTEx and a single-nucleus atlas indicates that the depot transcriptional architecture is reproducible, and single-nucleus data localize the ECM signal to stromal/progenitor rather than adipocyte compartments (Section 3.7); the BMI, CT, and histologic associations, however, remain specific to this cohort and require prospective replication. Planned extensions include a both-depot-trained fibrosis classifier with pathologist concordance, single-nucleus and spatial profiling of paired specimens [11, 25], and a longitudinal study relating baseline depot remodeling to post-surgical weight loss and metabolic response.

## 5. Conclusions

Human subcutaneous and omental VAT do not merely differ in location; they appear to implement complementary strategies for managing energy surplus. SAT preferentially engages a mesenchymal–ECM scaffold program consistent with high-capacity, long-term lipid storage, and structural containment, whereas omental VAT preferentially engages a mesothelial–stromal and vascular–hypoxia program consistent with a more dynamic, organ-adjacent depot that shows several-fold more adipose area per unit of fractional fibrosis. Depot-associated identity remained significant after adjustment for BMI and age, whereas the remodeling associations tracked adiposity, and histology revealed not only higher fractional fibrosis in SAT but also a different scaffold-to-storage relationship between the depots. Under chronic energy excess, these initially adaptive strategies may become maladaptive, producing SAT stiffness and restricted expandability on one side and VAT hypoxic vascular dysfunction on the other, without invoking distinct developmental lineages. Our findings provide a framework for future longitudinal studies to test whether depot-specific remodeling predicts metabolic outcomes or responses to weight loss interventions.

## Supporting information

Reproducibility Package (Source Code and Analysis Scripts)

Table S1 demographics and module definations

Table S2 paired SAT VAT DEG

Table S3 CT RNA mixed models

sample flow

storage vs scaffold

ThreeLayer validation

## List of Abbreviations

ASPCs: adipose stem/progenitor cells
BMI: body mass index
CD68: cluster of differentiation 68
CT: computed tomography
DEG: differentially expressed gene
DV200: percentage of RNA fragments >200 nucleotides
ECM: extracellular matrix
FDR: false discovery rate
GTEx: Genotype-Tissue Expression
HU: Hounsfield unit
IRB: Institutional Review Board
LOX: lysyl oxidase
PCA: principal component analysis
QC: quality control
RNA: ribonucleic acid
RNA-seq: RNA sequencing
SAT: subcutaneous adipose tissue
TMM: trimmed mean of M-values
VAT: visceral adipose tissue

## Data availability

Publicly available GTEx v8 adipose tissue gene expression data were obtained from the GTEx Portal. The Emont et al. human adipose single-nucleus dataset was obtained from the Single-Cell Portal (SCP1376) and Gene Expression Omnibus (GSE176171). The de-identified normalized expression matrix generated in the present study, analysis metadata, supplementary tables, and reproducibility scripts are deposited in Zenodo (DOI: 10.5281/zenodo.21421964).

## Author contributions: CRediT

G.O.K.: formal analysis, investigation, visualization, and writing of the original draft. D.H.K.: conceptualization, methodology, resources, supervision, project administration, funding acquisition, and writing (review and editing); corresponding author. S.-M.S.: investigation and validation (histopathology), resources, and writing (review and editing); co-corresponding author. Y.C.K.: investigation (physiological interpretation) and writing (review and editing). M.W.S.: resources and writing (review and editing). J.H.Y.: resources and writing (review and editing). All authors have reviewed and approved the final manuscript.

## Ethics

The Institutional Review Board of Chungbuk National University Hospital (IRB No. CBNU-2025-A-0174; approved 2025-12-29; expedited review). The IRB waived the requirement for informed consent for this retrospective analysis of archived specimens.

## Declaration of generative AI use

Declaration of generative AI use in the writing process. During the preparation of this paper, the authors used a generative AI tool to assist with language editing and formatting. The authors reviewed and edited the content as needed and take full responsibility for the content of the published articles.

