## Supplementary material for "Complementary remodeling strategies distinguish human subcutaneous and omental adipose tissue": Reproducibility Package (Source Code and Analysis Scripts): Figure6_external_validation.pdf

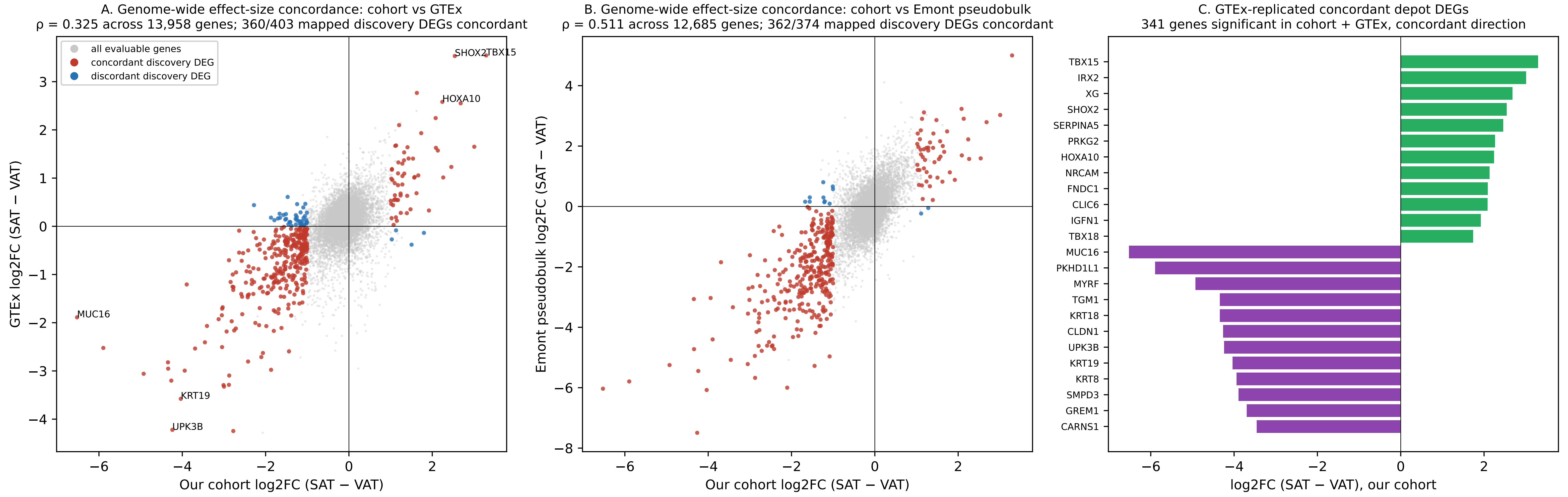


*Strong discovery DEGs: FDR < 0.05 and |log2FC| > 1 in our cohort. ρ computed over all shared genes; direction agreement over mapped strong discovery DEGs.*
