## Supplementary material for "Complementary remodeling strategies distinguish human subcutaneous and omental adipose tissue": Reproducibility Package (Source Code and Analysis Scripts): FigureS9_cellorigin.pdf

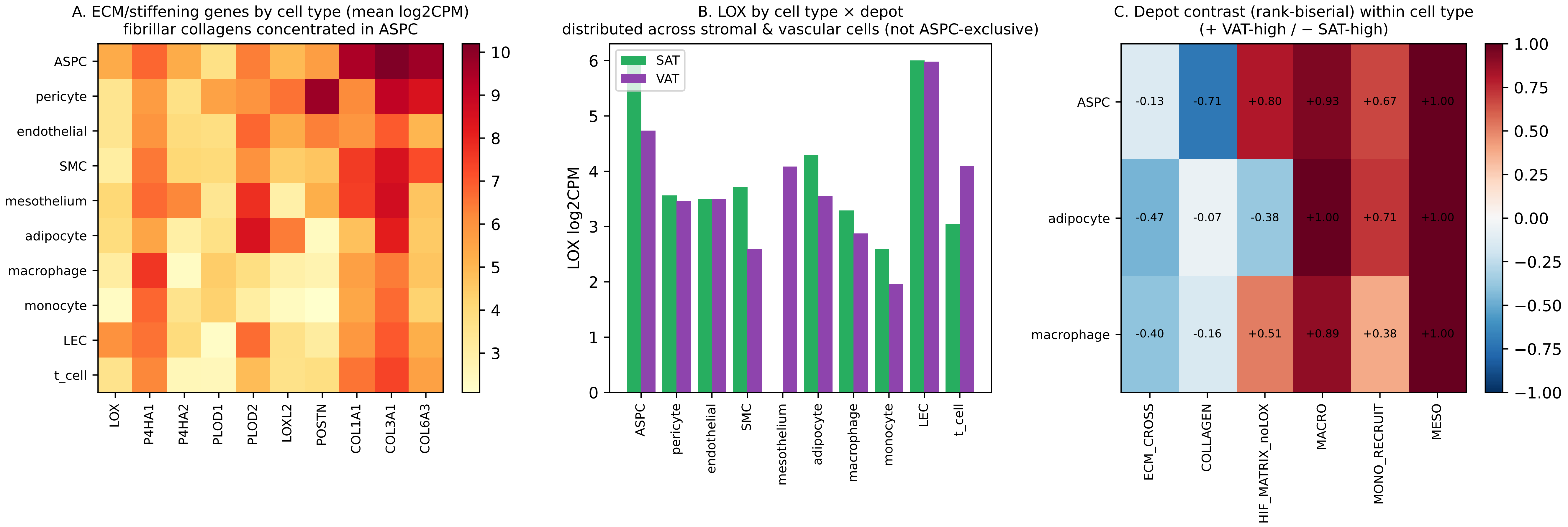


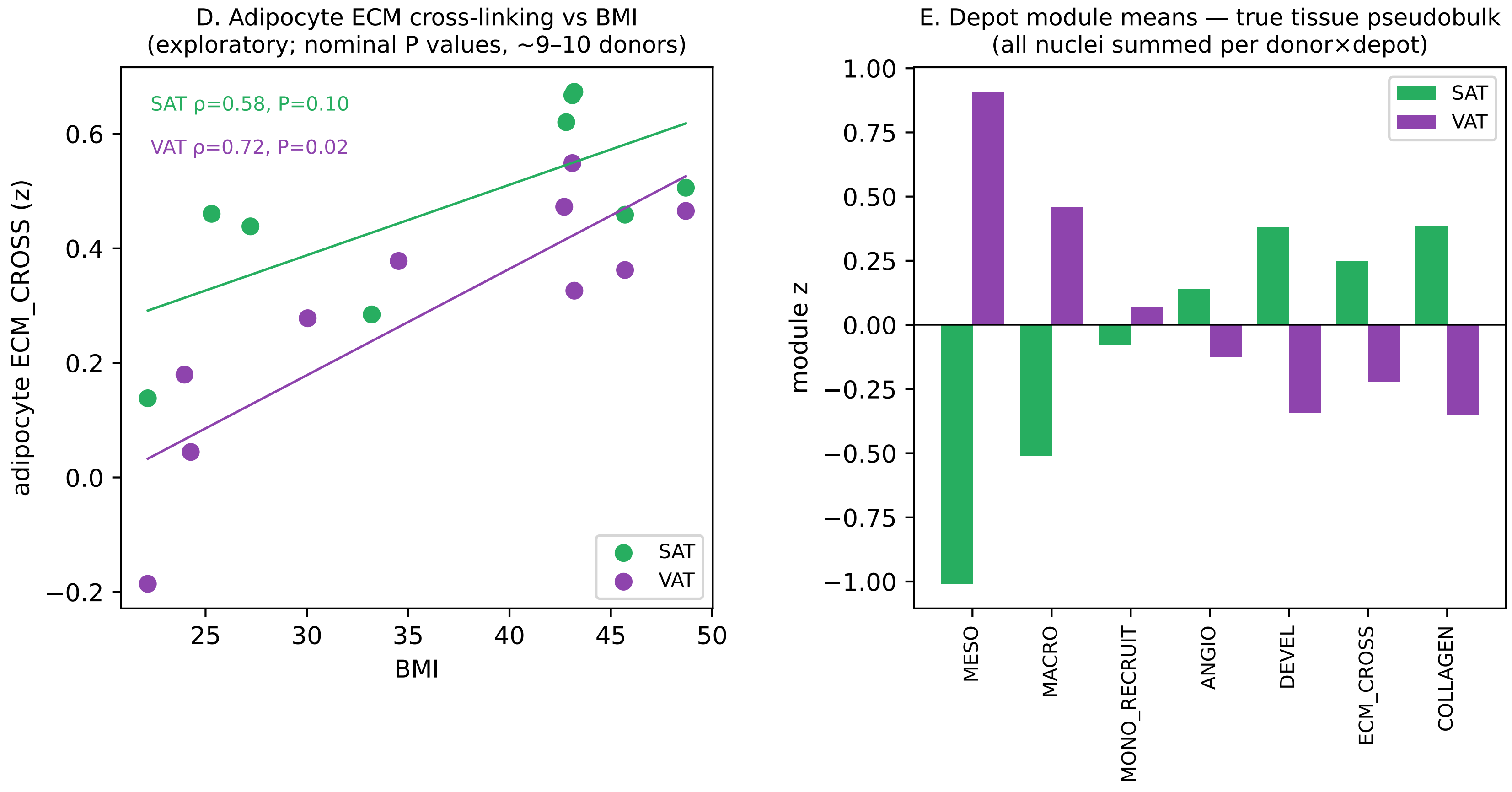


F. Cross-dataset concordance summary

| Comparison | Value |
|---|---|
| cohort–GTEx ρ (13,958 ge | 0.325 |
| discovery DEG concordan | 360/403 (89.3%) |
| cohort–Emont ρ (12,685 g | 0.511 |
| discovery DEG concordan | 362/374 (96.8%) |
| GTEx-replicated concorda | 341 |
| of these concordant in E | 308/317 (97.2%) |
