## Supplementary material for "Complementary remodeling strategies distinguish human subcutaneous and omental adipose tissue": Reproducibility Package (Source Code and Analysis Scripts): README.md

# Reproduction package — External validation of adipose depot transcriptional architecture

Manuscript: "Integrated transcriptomic, histologic, and CT analysis reveals distinct mesenchymal
and stromal remodeling programs in human subcutaneous and omental adipose tissue."

## What this reproduces
The three-tier external validation (our cohort n=29 -> GTEx n=1,204 -> Emont snRNA atlas):
genome-wide depot DEG concordance, the 341 GTEx-replicated concordant DEGs, cell-of-origin, and BMI analyses.

## Public data sources (download separately; not redistributed here)
- GTEx v8 adipose TPM (subcutaneous, visceral/omentum): GTEx Portal (gtexportal.org).
- Emont et al. 2022 human adipose single-nucleus atlas: Single Cell Portal SCP1376 and GEO GSE176171
  (files: Hs10X.counts.mtx.gz, Hs10X.counts.barcodes.tsv.gz, Hs10X.counts.features.tsv.gz, Hs.metadata.tsv).
- Our normalized expression matrix + metadata: to be deposited (Zenodo/GEO DOI before submission).

## scripts/
- GTEx_replication_20260712.py  : GTEx module-level replication (depot contrast, regulator~LOX).
- emont_gpu_pseudobulk.py        : GPU (CuPy) full-genome donor x depot x cell-type pseudobulk from the Emont mtx.
- Emont_pseudobulk_build.py      : CPU streaming pseudobulk (module genes).
- mechanism_analysis.py          : original n=29 module-score / mechanism analysis.

## data/ (processed outputs)
- OurCohort_genomewide_depot_DEG.csv, GTEx_genomewide_depot_DEG.csv, Emont_genomewide_depot_DEG_paired.csv
- ThreeLayer_DEG_merged.csv       : per-gene log2FC/FDR in all three datasets +
                                    concordance flags. Columns: concordant_ours_gtex
                                    (sign(ours)==sign(GTEx)) and concordant_3way_true
                                    (sign agrees across ours, GTEx AND Emont). These match
                                    SuppTable_ThreeLayer_validation.xlsx exactly.
- Emont_FULL_pseudobulk_*.csv     : 29,093 genes x 273 donor x depot x cell-type pseudobulks.
- SuppTable_ThreeLayer_validation.xlsx : Summary, column definitions, 341-DEG list, all-gene 3-layer table.

## Key numbers (verifiable from data/)
- cohort~GTEx: Spearman rho=0.325 (13,958 genes); 360/403 strong discovery DEGs concordant (89.3%).
- cohort~Emont: rho=0.511 (12,685 genes); 362/374 concordant (96.8%).
- 341 GTEx-replicated concordant DEGs; of 317 with Emont, 308 concordant (97.2%).
- Strong discovery DEGs: our-cohort Welch t-test, FDR<0.05 & |log2FC|>1 (420 genes).

## Dependencies
python3, numpy, pandas, scipy, statsmodels, matplotlib; cupy-cuda12x optional (GPU).

## How to run
```bash
python -m pip install -r requirements.txt

# Step 0 — verify the headline numbers directly from shipped data/ (no downloads):
python run_all.py --verify

# Full regeneration from raw inputs (download separately; see sources above):
python run_all.py \
    --cohort-dir /path/to/cohort \    # norm.count.txt + CT-RNA xlsx (Zenodo/GEO DOI)
    --emont-dir  /path/to/emont  \    # Hs10X.counts.* + Hs.metadata.tsv (SCP1376/GSE176171)
    --gtex-dir   /path/to/gtex   \    # GTEx v8 adipose .gct.gz (GTEx Portal)
    --out        outputs
```
All scripts also accept the same flags individually, or the environment variables
`COHORT_DIR`, `EMONT_DIR`, `GTEX_DIR`, `OUT_DIR`. No paths are hardcoded.

## Reproduction scope (what runs with what)
- **Without any download:** `run_all.py --verify` recomputes the three-tier concordance
  statistics (cohort~GTEx rho, cohort~Emont rho, the ours/GTEx and true 3-way DEG counts)
  directly from `data/`. This is the package's self-contained check.
- **With raw inputs:** the four scripts regenerate the processed tables/pseudobulks and
  the depot-DEG contrasts. Supplementary Figures S1/S2/S7 are regenerated by
  `scripts/make_supp_figures.py` once the cohort `norm.count.txt` + metadata are available
  (deposited on Zenodo/GEO before submission).
- The BMI-, CT-, and histology-specific analyses depend on the cohort matrix and are
  reproducible once that matrix is deposited; the public-dataset architecture (GTEx/Emont)
  is reproducible now from the shipped tables.

## Integrity
`MANIFEST_SHA256.txt` lists a SHA-256 for every shipped file. Verify with:
```bash
sha256sum -c MANIFEST_SHA256.txt
```
