## Supplementary material for "Complementary remodeling strategies distinguish human subcutaneous and omental adipose tissue": Reproducibility Package (Source Code and Analysis Scripts): requirements.txt

# Reproduction package — Python dependencies
# Tested with Python 3.12
numpy>=1.26
pandas>=2.0
scipy>=1.11
statsmodels>=0.14
scikit-learn>=1.3
matplotlib>=3.7
openpyxl>=3.1
# Optional, only for the GPU pseudobulk path (emont_gpu_pseudobulk.py):
# cupy-cuda12x   # install matching your CUDA toolkit; CPU fallback is automatic
