## Supplementary material for "Complementary remodeling strategies distinguish human subcutaneous and omental adipose tissue": Reproducibility Package (Source Code and Analysis Scripts): run_all.py

```
#!/usr/bin/env python3
"""
run_all.py — driver for the external-validation reproduction package.

This package reproduces the THREE-TIER EXTERNAL VALIDATION
(our cohort n=29  ->  GTEx n=1,204  ->  Emont snRNA atlas):
genome-wide depot-DEG concordance, the 341 GTEx-replicated concordant DEGs,
cell-of-origin, and the BMI analyses.

Two kinds of inputs are involved:
  (1) PROCESSED outputs already shipped in data/ — every "Key number" in the
      README is directly verifiable from these without re-downloading anything
      (see step 0 below).
  (2) RAW public/cohort inputs (NOT redistributed) — needed only to regenerate
      the processed outputs from scratch. Point the scripts at them via flags:
        --gtex-dir   GTEx v8 adipose .gct.gz files      (GTEx Portal)
        --emont-dir  Hs10X.counts.* + Hs.metadata.tsv   (SCP1376 / GSE176171)
        --cohort-dir norm.count.txt + CT-RNA xlsx        (Zenodo/GEO DOI, on deposit)

Usage:
  python run_all.py --verify                # step 0 only: recompute key numbers from data/
  python run_all.py --emont-dir DIR --gtex-dir DIR --cohort-dir DIR --out outputs
"""
import argparse, os, subprocess, sys, hashlib, json

HERE = os.path.dirname(os.path.abspath(__file__))

def sha256(path, buf=1<<20):
    h = hashlib.sha256()
    with open(path, "rb") as f:
        for chunk in iter(lambda: f.read(buf), b""):
            h.update(chunk)
    return h.hexdigest()

def verify_key_numbers():
    """Recompute the headline concordance stats directly from data/ThreeLayer_DEG_merged.csv
    and the shipped genome-wide DEG tables. No raw downloads needed."""
    import pandas as pd, numpy as np
    d = os.path.join(HERE, "data")
    tl = pd.read_csv(os.path.join(d, "ThreeLayer_DEG_merged.csv"), index_col=0)
    print("[verify] ThreeLayer_DEG_merged.csv rows:", len(tl))
    print("[verify] concordant_ours_gtex :", int(tl['concordant_ours_gtex'].sum()), "/", len(tl))
    print("[verify] concordant_3way_true :", int(tl['concordant_3way_true'].sum()), "/", len(tl))
    # cohort~GTEx / cohort~Emont Spearman over shared genes
    from scipy.stats import spearmanr
    m_og = tl.dropna(subset=['LFC_ours','LFC_gtex'])
    rho_og = spearmanr(m_og['LFC_ours'], m_og['LFC_gtex']).correlation
    m_oe = tl.dropna(subset=['LFC_ours','LFC_emont'])
    rho_oe = spearmanr(m_oe['LFC_ours'], m_oe['LFC_emont']).correlation
    print(f"[verify] cohort~GTEx  Spearman rho = {rho_og:.3f}  (n={len(m_og)} shared genes)")
    print(f"[verify] cohort~Emont Spearman rho = {rho_oe:.3f}  (n={len(m_oe)} shared genes)")
    print("[verify] (Manuscript reports rho_GTEx=0.325, rho_Emont=0.511 on the "
          "reported gene subsets; see README for the exact gene-set definitions.)")

def run(cmd):
    print("[run]", " ".join(cmd)); sys.stdout.flush()
    subprocess.run(cmd, check=True)

def main():
    ap = argparse.ArgumentParser()
    ap.add_argument("--verify", action="store_true",
                    help="only recompute key numbers from shipped data/ (no raw inputs needed)")
    ap.add_argument("--gtex-dir"); ap.add_argument("--emont-dir"); ap.add_argument("--cohort-dir")
    ap.add_argument("--out", default=os.path.join(HERE, "outputs"))
    a = ap.parse_args()
    os.makedirs(a.out, exist_ok=True)

    # step 0: always available
    verify_key_numbers()
    if a.verify:
        return

    S = os.path.join(HERE, "scripts")
    if a.cohort_dir:
        run([sys.executable, os.path.join(S, "mechanism_analysis.py"),
             "--cohort-dir", a.cohort_dir, "--out", a.out])
    if a.emont_dir:
        run([sys.executable, os.path.join(S, "Emont_pseudobulk_build.py"),
             "--emont-dir", a.emont_dir, "--out", a.out])
    if a.gtex_dir:
        run([sys.executable, os.path.join(S, "GTEx_replication_20260712.py"),
             "--gtex-dir", a.gtex_dir, "--out", a.out])
    print("\n[done] Provide --cohort-dir/--emont-dir/--gtex-dir to regenerate the "
          "corresponding processed outputs. For the Supplementary Figures S1/S2/S7 "
          "regeneration, see scripts/make_supp_figures.py (needs --cohort-dir).")

if __name__ == "__main__":
    main()
```
