## Supplementary material for "Complementary remodeling strategies distinguish human subcutaneous and omental adipose tissue": Reproducibility Package (Source Code and Analysis Scripts): emont_gpu_pseudobulk.py

```
#!/usr/bin/env python3
# -*- coding: utf-8 -*-
"""
Emont SCP1376 human snRNA-seq  ->  FULL-genome donor x depot x cell-type pseudobulk (GPU).
- 29,093 genes x 137,684 nuclei -> aggregate ALL genes by (donor x depot x cell-type)
- GPU (CuPy) sparse matmul; auto CPU fallback if CuPy not installed
- outputs the validated RAW pseudobulk matrix + a validation summary
Owner: 김대훈 | run on RTX 4090 (CUDA 12)
Files expected in the SAME folder as this script:
  Hs10X.counts.mtx.gz, Hs10X.counts.barcodes.tsv.gz, Hs10X.counts.features.tsv.gz, Hs.metadata.tsv
"""
import os, sys, gzip, time, json
import numpy as np, pandas as pd
from scipy import stats
import scipy.sparse as sp

HERE = os.path.dirname(os.path.abspath(__file__))
def P(name): return os.path.join(HERE, name)
MTX, BC, FEAT, MD = P("Hs10X.counts.mtx.gz"), P("Hs10X.counts.barcodes.tsv.gz"), P("Hs10X.counts.features.tsv.gz"), P("Hs.metadata.tsv")
OUTDIR = P("Emont_FULL_output"); os.makedirs(OUTDIR, exist_ok=True)
t0 = time.time()

# ---------- GPU or CPU ----------
USE_GPU = False
try:
    import cupy as cp
    import cupyx.scipy.sparse as csp
    cp.cuda.runtime.getDeviceCount()
    USE_GPU = True
    print(f"[GPU] CuPy detected -> {cp.cuda.runtime.getDeviceProperties(0)['name'].decode()}")
except Exception as e:
    print(f"[CPU] CuPy not usable ({e}); falling back to CPU scipy.")

ALIAS = {'CYR61':'CCN1','CTGF':'CCN2'}

# ---------- 1. features & barcodes ----------
print("[1/5] loading features / barcodes / metadata ...", flush=True)
feats = [l.rstrip('\n').split('\t') for l in gzip.open(FEAT,'rt')]
gene_sym = [ALIAS.get(f[1], f[1]) for f in feats]            # symbol per gene row (0-based)
nGenes = len(gene_sym)
barcodes = [l.strip() for l in gzip.open(BC,'rt')]
nCells = len(barcodes)

md = pd.read_csv(MD, sep='\t', low_memory=False).set_index('cell_id').loc[barcodes].reset_index()
md['ctype'] = md['cell_type__custom'].astype(str)
md['depot'] = md['fat__type'].astype(str)
md['grp']   = md['donor_id'] + '|' + md['depot'] + '|' + md['ctype']
cats = md['grp'].astype('category')
gid  = cats.cat.codes.values.astype(np.int32)               # 0-based group id per cell
groups = list(cats.cat.categories)
nGroups = len(groups)
print(f"    genes={nGenes}  cells={nCells}  groups={nGroups}")

# ---------- 2. read the matrix triplets ----------
print("[2/5] reading count matrix (this reads ~257M nonzeros) ...", flush=True)
# MatrixMarket: line1 %%..., line2 'nGenes nCells nnz', then g c v (1-based)
arr = pd.read_csv(MTX, sep=' ', skiprows=2, header=None, dtype=np.int32).values
g = arr[:,0]-1; c = arr[:,1]-1; v = arr[:,2].astype(np.float32)
del arr
print(f"    nnz={len(v):,}  ({time.time()-t0:.0f}s)")

# ---------- 3. aggregate genes x groups (GPU) ----------
print(f"[3/5] aggregating on {'GPU' if USE_GPU else 'CPU'} ...", flush=True)
# counts: genes x cells  (CSR)   ;   onehot: cells x groups (CSR)  ;  pb = counts @ onehot
if USE_GPU:
    counts = csp.csr_matrix((cp.asarray(v), (cp.asarray(g), cp.asarray(c))), shape=(nGenes, nCells))
    onehot = csp.csr_matrix((cp.ones(nCells, dtype=cp.float32), (cp.arange(nCells), cp.asarray(gid))),
                            shape=(nCells, nGroups))
    pb = (counts @ onehot).toarray()          # genes x groups (dense)
    pb = cp.asnumpy(pb)
else:
    counts = sp.csr_matrix((v, (g, c)), shape=(nGenes, nCells))
    onehot = sp.csr_matrix((np.ones(nCells, np.float32), (np.arange(nCells), gid)), shape=(nCells, nGroups))
    pb = (counts @ onehot).toarray()
del counts, onehot, g, c, v
print(f"    pseudobulk counts matrix = {pb.shape}  ({time.time()-t0:.0f}s)")

# ---------- 4. CPM + log2, save RAW ----------
print("[4/5] normalizing + saving RAW pseudobulk ...", flush=True)
libsize = pb.sum(0)                            # total UMIs per group
cpm = pb / np.where(libsize==0,1,libsize) * 1e6
log2cpm = np.log2(cpm + 1)

gmeta = md.groupby('grp').agg(
    donor=('donor_id','first'), depot=('depot','first'), ctype=('ctype','first'),
    n_cells=('grp','size'),
    bmi=('bmi', lambda s: pd.to_numeric(s,errors='coerce').mean()),
    bmi_group=('bmi__group', lambda s: s.mode().iat[0])).reindex(groups).reset_index()
gmeta['libsize'] = libsize

genes_idx = pd.Index(gene_sym, name='gene')
raw_counts_df = pd.DataFrame(pb.astype(np.int64), index=genes_idx, columns=groups)
log2_df       = pd.DataFrame(log2cpm.astype(np.float32), index=genes_idx, columns=groups)
# collapse duplicate gene symbols -> keep max-mean row
log2_df['m'] = log2_df.mean(1); log2_df = log2_df.sort_values('m',ascending=False)[~log2_df.index.duplicated()].drop(columns='m')
raw_counts_df = raw_counts_df.loc[log2_df.index]

raw_counts_df.to_csv(os.path.join(OUTDIR,"Emont_FULL_pseudobulk_counts.csv"))
log2_df.to_csv(os.path.join(OUTDIR,"Emont_FULL_pseudobulk_log2cpm.csv"))
gmeta.to_csv(os.path.join(OUTDIR,"Emont_FULL_pseudobulk_meta.csv"), index=False)
print(f"    saved RAW: {raw_counts_df.shape[0]} genes x {raw_counts_df.shape[1]} pseudobulk samples")

# ---------- 5. validation summary ----------
print("[5/5] validation summary ...", flush=True)
keep = gmeta['n_cells'].values >= 10
L = log2_df.loc[:, gmeta['grp'][keep]]
GM = gmeta[keep].reset_index(drop=True); GM.index = L.columns
Z = L.sub(L.mean(1),axis=0).div(L.std(1,ddof=1).replace(0,np.nan),axis=0)
modules = {
 'HIF_MATRIX_noLOX':['P4HA1','P4HA2','PLOD1','PLOD2','LOXL2'],
 'ECM_CROSS':['LOX','LOXL1','LOXL2','POSTN','P4HA1','PLOD2'],
 'COLLAGEN':['COL1A1','COL1A2','COL3A1','COL6A3'],
 'MACRO':['CD68','CD163','MRC1','MSR1','CSF1R','ITGAM','AIF1','TYROBP','C1QA','C1QB'],
 'MONO_RECRUIT':['CCL2','CCR2','CCL7','CCL8','CX3CR1','S100A8','S100A9'],
 'MESO':['UPK3B','KRT19'],'DEVEL':['TBX15','SHOX2','HOXA5','HOXA10','NR2F1']}
def modscore(gs):
    gg=[x for x in gs if x in Z.index]; return Z.loc[gg].mean(0)
S = pd.DataFrame({k:modscore(v) for k,v in modules.items()}); S['depot']=GM['depot'].values; S['ctype']=GM['ctype'].values; S['bmi']=GM['bmi'].values
rows=[]
for ct in ['ASPC','adipocyte','macrophage']:
    sub=S[S.ctype==ct]
    for m in modules:
        a=sub.loc[sub.depot=='SAT',m].dropna(); b=sub.loc[sub.depot=='VAT',m].dropna()
        if len(a)>=3 and len(b)>=3:
            u,p=stats.mannwhitneyu(a,b); rb=1-2*u/(len(a)*len(b))
            rows.append(dict(ctype=ct,module=m,SAT=round(a.mean(),3),VAT=round(b.mean(),3),
                             dir='VAT>SAT' if b.mean()>a.mean() else 'SAT>VAT',rank_biserial=round(rb,3),p=round(p,4)))
pd.DataFrame(rows).to_csv(os.path.join(OUTDIR,"Emont_FULL_validation_depot_contrast.csv"),index=False)
# cell-of-origin for stiffening genes
stiff=[x for x in ['LOX','P4HA1','PLOD2','LOXL2','POSTN','COL1A1','COL3A1','COL6A3'] if x in log2_df.index]
byct=log2_df.loc[stiff].T.join(gmeta.set_index('grp')['ctype']).groupby('ctype').mean()
byct.round(2).to_csv(os.path.join(OUTDIR,"Emont_FULL_celltype_stiffening_expr.csv"))

print("\n===== VALIDATION SNAPSHOT =====")
print("LOX by cell type (top):")
print(byct['LOX'].sort_values(ascending=False).head(5).round(2).to_string())
print("\nDepot contrast (ASPC):")
for r in rows:
    if r['ctype']=='ASPC': print(f"  {r['module']:16s} {r['dir']:>8s} rb={r['rank_biserial']:+.3f} p={r['p']}")
print(f"\nALL DONE in {time.time()-t0:.0f}s. Outputs in: {OUTDIR}")
json.dump({'genes':int(raw_counts_df.shape[0]),'pseudobulk_samples':int(raw_counts_df.shape[1]),
           'used_gpu':USE_GPU,'seconds':round(time.time()-t0,1)}, open(os.path.join(OUTDIR,"run_info.json"),"w"), indent=1)
```
