## Supplementary material for "Complementary remodeling strategies distinguish human subcutaneous and omental adipose tissue": Reproducibility Package (Source Code and Analysis Scripts): Emont_pseudobulk_build.py

```
#!/usr/bin/env python3
"""Emont SCP1376 human snRNA-seq -> donor x depot x cell-type pseudobulk.
Streams the 257M-nonzero MatrixMarket file, keeping only target module genes,
and library size per cell for CPM. NO cell-as-sample (donor pseudobulk only)."""
import gzip, numpy as np, pandas as pd, json, warnings
warnings.filterwarnings('ignore')
import argparse, os
_ap = argparse.ArgumentParser(description="Emont snRNA -> pseudobulk (CPU streaming)")
_ap.add_argument("--emont-dir", default=os.environ.get("EMONT_DIR", "./emont"),
                 help="directory holding Hs10X.counts.* and Hs.metadata.tsv")
_ap.add_argument("--out", default=os.environ.get("OUT_DIR", "./outputs"))
_args, _ = _ap.parse_known_args()
BASE = _args.emont_dir.rstrip("/") + "/"
OUT = _args.out.rstrip("/") + "/"
os.makedirs(OUT, exist_ok=True)
MTX=BASE+"Hs10X.counts.mtx.gz"; FEAT=BASE+"Hs10X.counts.features.tsv.gz"; BC=BASE+"Hs10X.counts.barcodes.tsv.gz"; MD=BASE+"Hs.metadata.tsv"

# ---- module gene sets (same as GTEx, with alias handling) ----
ALIAS={'CYR61':'CCN1','CTGF':'CCN2'}
modules={
'HYPOXIA':['VEGFA','SLC2A1','LDHA','PGK1','ADM','PDK1','BNIP3','ANGPTL4','EGLN3','NDRG1','ALDOA','ENO1','HK2','PFKFB3'],
'HIF_MATRIX':['P4HA1','P4HA2','PLOD1','PLOD2','LOX','LOXL2'],
'HIF_MATRIX_noLOX':['P4HA1','P4HA2','PLOD1','PLOD2','LOXL2'],
'TGFB':['SERPINE1','SMAD7','TGFBI','SKIL','JUNB','LTBP2','TGFB1','TGFBR1','THBS1','CCN2'],
'YAPTAZ':['CCN2','CCN1','AMOTL2','AXL','THBS1','F3','TAGLN','ACTA2','MYL9','TEAD1'],
'ECM_CROSS':['LOX','LOXL1','LOXL2','POSTN','P4HA1','PLOD2'],
'COLLAGEN':['COL1A1','COL1A2','COL3A1','COL6A3'],
'MACRO':['CD68','CD163','MRC1','MSR1','CSF1R','ITGAM','AIF1','TYROBP','C1QA','C1QB'],
'MONO_RECRUIT':['CCL2','CCR2','CCL7','CCL8','CX3CR1','S100A8','S100A9'],
'ANGIO':['VEGFA','KDR','FLT1','PECAM1','NRP2','APLNR','CCN1','ESM1'],
'DEVEL':['TBX15','SHOX2','HOXA5','HOXA10','NR2F1'],'MESO':['UPK3B','KRT19']}
EXPR=['LOX','LOXL1','LOXL2','P4HA1','P4HA2','PLOD1','PLOD2','POSTN','COL1A1','COL3A1','COL6A3',
      'HIF1A','EPAS1','TGFB1','WWTR1','YAP1','CD68','CCL2','UPK3B']
targets=set(EXPR)
for gs in modules.values(): targets|=set(gs)

# ---- features: row(1-based) -> symbol (apply alias) ----
feats=[l.rstrip('\n').split('\t') for l in gzip.open(FEAT,'rt')]
sym=[ALIAS.get(f[1],f[1]) for f in feats]           # gene symbol per row (0-based)
row_of={}                                            # local target index
tgt_syms=[]
for i,s in enumerate(sym):
    if s in targets and s not in row_of:
        row_of[i]=len(tgt_syms); tgt_syms.append(s)  # map mtx row(0-based) -> local
present=set(tgt_syms); print("target genes found:",len(tgt_syms),"/",len(targets),"| missing:",sorted(targets-present))
nT=len(tgt_syms); nCells=len(open(BC:=BC).readlines()) if False else None
# barcodes + metadata -> per-cell group
bc=[l.strip() for l in gzip.open(BASE+"Hs10X.counts.barcodes.tsv.gz",'rt')]
nCells=len(bc)
md=pd.read_csv(MD,sep='\t',low_memory=False).set_index('cell_id')
md=md.loc[bc]  # align to matrix column order
donor=md['donor_id'].values; depot=md['fat__type'].values; ctype=md['cell_type__custom'].astype(str).values
bmi=pd.to_numeric(md['bmi'],errors='coerce').values; bmigrp=md['bmi__group'].astype(str).values

# ---- stream mtx, accumulate ----
tgt_by_cell=np.zeros((nT,nCells),dtype=np.float64)
cell_total=np.zeros(nCells,dtype=np.float64)
row_is_tgt=np.zeros(len(sym),dtype=np.int32)-1
for r,loc in row_of.items(): row_is_tgt[r]=loc
CH=25_000_000
it=pd.read_csv(MTX,sep=' ',skiprows=2,header=None,names=['g','c','v'],dtype=np.int32,chunksize=CH)
tot=0
for ch in it:
    g=ch['g'].values-1; c=ch['c'].values-1; v=ch['v'].values.astype(np.float64)
    np.add.at(cell_total,c,v)                       # library size (all genes)
    loc=row_is_tgt[g]; keep=loc>=0
    if keep.any():
        np.add.at(tgt_by_cell,(loc[keep],c[keep]),v[keep])
    tot+=len(ch); print("  processed",tot,"nnz",flush=True)

# ---- collapse to donor x depot x celltype pseudobulk ----
grp=pd.DataFrame({'donor':donor,'depot':depot,'ctype':ctype,'idx':np.arange(nCells)})
rows=[]; meta_rows=[]
GT=pd.Series(cell_total)
for (d,dep,ct),sub in grp.groupby(['donor','depot','ctype']):
    ix=sub['idx'].values
    ncell=len(ix); libtot=cell_total[ix].sum()
    if ncell<10 or libtot<=0: continue              # min 10 nuclei per pseudobulk
    genecnt=tgt_by_cell[:,ix].sum(1)
    cpm=genecnt/libtot*1e6
    rows.append(np.log2(cpm+1))
    b=bmi[ix]; meta_rows.append(dict(donor=d,depot=dep,ctype=ct,n_cells=ncell,
        bmi=float(np.nanmean(b)),bmi_group=pd.Series(bmigrp[ix]).mode().iat[0]))
PB=pd.DataFrame(rows,columns=tgt_syms); M=pd.DataFrame(meta_rows)
PB.index=M.index
print("\npseudobulk samples (donor x depot x celltype, >=10 cells):",PB.shape[0])
print(M.groupby(['depot','ctype']).size().unstack(fill_value=0))

PB.to_csv(OUT+"Emont_pseudobulk_log2cpm.csv"); M.to_csv(OUT+"Emont_pseudobulk_meta.csv",index=False)
# also save a genes/modules json for downstream
json.dump({'modules':modules,'targets':tgt_syms,'missing':sorted(targets-present)},open(OUT+"Emont_modules.json","w"),indent=1)
print("SAVED pseudobulk")
```
