## Supplementary material for "Complementary remodeling strategies distinguish human subcutaneous and omental adipose tissue": Reproducibility Package (Source Code and Analysis Scripts): GTEx_replication_20260712.py

```
#!/usr/bin/env python3
"""GTEx v8 large-cohort replication of the n=29 depot-remodeling findings.
Uses the EXACT module definitions from mechanism_analysis.py.
SAT = adipose subcutaneous, VAT = adipose visceral (omentum).
Public GTEx TPM has no BMI -> BMI-dependent claims are NOT tested here."""
import pandas as pd, numpy as np, json, gzip, warnings
from scipy import stats
warnings.filterwarnings('ignore'); np.random.seed(1)

import argparse, os
_ap = argparse.ArgumentParser(description="GTEx replication of depot findings")
_ap.add_argument("--gtex-dir", default=os.environ.get("GTEX_DIR", "./gtex"),
                 help="directory holding the two GTEx adipose .gct.gz files")
_ap.add_argument("--out", default=os.environ.get("OUT_DIR", "./outputs"),
                 help="output directory")
_args, _ = _ap.parse_known_args()
UP = _args.gtex_dir.rstrip("/") + "/"
OUT = _args.out.rstrip("/") + "/"
os.makedirs(OUT, exist_ok=True)
FILES={'SAT':UP+"gene_tpm_2017-06-05_v8_adipose_subcutaneous.gct.gz",
       'VAT':UP+"gene_tpm_2017-06-05_v8_adipose_visceral_omentum.gct.gz"}

# ---- exact module definitions (from mechanism_analysis.py) ----
HYPOXIA=['VEGFA','SLC2A1','LDHA','PGK1','ADM','PDK1','BNIP3','ANGPTL4','EGLN3','NDRG1','ALDOA','ENO1','HK2','PFKFB3']
HIF_MATRIX=['P4HA1','P4HA2','PLOD1','PLOD2','LOX','LOXL2']
TGFB=['SERPINE1','SMAD7','TGFBI','SKIL','JUNB','LTBP2','TGFB1','TGFBR1','THBS1','CCN2']
YAPTAZ=['CCN2','CCN1','AMOTL2','AXL','THBS1','F3','TAGLN','ACTA2','MYL9','TEAD1']
ECM_CROSS=['LOX','LOXL1','LOXL2','POSTN','P4HA1','PLOD2']
COLLAGEN=['COL1A1','COL1A2','COL3A1','COL6A3']
MACRO=['CD68','CD163','MRC1','MSR1','CSF1R','ITGAM','AIF1','TYROBP','C1QA','C1QB']
MONO_RECRUIT=['CCL2','CCR2','CCL7','CCL8','CX3CR1','S100A8','S100A9']
ANGIO=['VEGFA','KDR','FLT1','PECAM1','NRP2','APLNR','CCN1','ESM1']
DEVEL=['TBX15','SHOX2','HOXA5','HOXA10','NR2F1']; MESO=['UPK3B','KRT19']
modules={'HYPOXIA':HYPOXIA,'HIF_MATRIX':HIF_MATRIX,'TGFB':TGFB,'YAPTAZ':YAPTAZ,'ECM_CROSS':ECM_CROSS,
 'COLLAGEN':COLLAGEN,'MACRO':MACRO,'MONO_RECRUIT':MONO_RECRUIT,'ANGIO':ANGIO,'DEVEL':DEVEL,'MESO':MESO}
EXPR=['LOX','POSTN','LOXL2','COL1A1','COL3A1','COL6A3','HIF1A','EPAS1','TGFB1','WWTR1','YAP1','PIEZO1','PTK2','CCN2','CD68','CD163']
need=set(EXPR)
for gs in modules.values(): need|=set(gs)

# ---- load only needed gene rows from each GCT ----
def load_gct(path):
    # gct: line1 version, line2 dims, line3 header (id Name Description samples...)
    df=pd.read_csv(path,sep='\t',skiprows=2,low_memory=False)
    # column 'Description' = gene symbol; drop id/Name
    keep=df[df['Description'].isin(need)].copy()
    samp=[c for c in df.columns if c.startswith('GTEX-')]
    keep=keep[['Description']+samp]
    # collapse duplicate symbols -> max mean expression row
    keep['m']=keep[samp].mean(1)
    keep=keep.sort_values('m',ascending=False).drop_duplicates('Description').drop(columns='m')
    return keep.set_index('Description')[samp]

mats={}
for depot,p in FILES.items():
    mats[depot]=load_gct(p)
    print(f"{depot}: {mats[depot].shape[1]} samples, {mats[depot].shape[0]} of {len(need)} target genes found")

# align genes, combine
genes=sorted(set(mats['SAT'].index)|set(mats['VAT'].index))
tpm=pd.concat([mats['SAT'].reindex(genes),mats['VAT'].reindex(genes)],axis=1)
depot_lab=pd.Series(['SAT']*mats['SAT'].shape[1]+['VAT']*mats['VAT'].shape[1],index=tpm.columns)
SAT=depot_lab.index[depot_lab=='SAT'].tolist(); VAT=depot_lab.index[depot_lab=='VAT'].tolist()
print(f"combined: SAT={len(SAT)} VAT={len(VAT)} total={tpm.shape[1]}")
missing=need-set(genes)
if missing: print("MISSING genes (not in GTEx):",sorted(missing))

log2=np.log2(tpm+1)
Z=log2.sub(log2.mean(1),axis=0).div(log2.std(1,ddof=1).replace(0,np.nan),axis=0)
cols=tpm.columns.tolist()
def modz(gs):
    gg=[g for g in gs if g in Z.index]; return Z.loc[gg].mean(0),gg
S={}; used={}
for k,gs in modules.items(): S[k],used[k]=modz(gs)
S=pd.DataFrame(S); S['depot']=depot_lab
for g in EXPR:
    if g in log2.index: S[g+'_expr']=log2.loc[g]

def spear(a,b):
    m=(~pd.isna(a))&(~pd.isna(b)); n=int(m.sum())
    if n<4: return (np.nan,np.nan,n)
    rho,p=stats.spearmanr(a[m],b[m]); return (round(float(rho),3),float(p),n)

results={'cohort':{'SAT':len(SAT),'VAT':len(VAT)},'genes_used':used,'missing':sorted(missing)}

# ---- A. depot contrast (rank-biserial) ----
contrast=[]
for k in ['HYPOXIA','HIF_MATRIX','ECM_CROSS','COLLAGEN','MACRO','MONO_RECRUIT','TGFB','YAPTAZ','ANGIO','DEVEL','MESO','LOX_expr','POSTN_expr','CD68_expr']:
    if k not in S: continue
    a=S.loc[SAT,k].dropna(); b=S.loc[VAT,k].dropna(); u,p=stats.mannwhitneyu(a,b,alternative='two-sided')
    rb=1-2*u/(len(a)*len(b))
    contrast.append(dict(module=k,SAT_mean=round(float(a.mean()),3),VAT_mean=round(float(b.mean()),3),
        dir='SAT>VAT' if a.mean()>b.mean() else 'VAT>SAT',rank_biserial=round(float(rb),3),p=p))
results['depot_contrast']=contrast

# ---- B. upstream regulator -> LOX / POSTN, within depot ----
reg_rows=[]
for depot,cc in [('SAT',SAT),('VAT',VAT)]:
    sub=S.loc[cc]
    for reg in ['HYPOXIA','TGFB','YAPTAZ','HIF_MATRIX']:
        for outc in ['LOX_expr','POSTN_expr']:
            if outc in sub:
                rho,p,n=spear(sub[reg],sub[outc]); reg_rows.append(dict(depot=depot,regulon=reg,outcome=outc,rho=rho,p=p,n=n))
    for tf in ['HIF1A_expr','EPAS1_expr','TGFB1_expr','WWTR1_expr','YAP1_expr','PIEZO1_expr','PTK2_expr']:
        if tf in sub: rho,p,n=spear(sub[tf],sub['LOX_expr']); reg_rows.append(dict(depot=depot,regulon=tf,outcome='LOX_expr',rho=rho,p=p,n=n))
results['upstream_regulator']=reg_rows

# ---- C. network dichotomy ----
net={}
for depot,cc in [('SAT',SAT),('VAT',VAT)]:
    sub=S.loc[cc]
    net[depot]={'hypoxia~ECMcross':spear(sub['HYPOXIA'],sub['ECM_CROSS']),
        'hypoxia~macro':spear(sub['HYPOXIA'],sub['MACRO']),'ECMcross~macro':spear(sub['ECM_CROSS'],sub['MACRO']),
        'macro~mono_recruit':spear(sub['MACRO'],sub['MONO_RECRUIT']),'ECMcross~TGFB':spear(sub['ECM_CROSS'],sub['TGFB']),
        'ECMcross~YAPTAZ':spear(sub['ECM_CROSS'],sub['YAPTAZ'])}
results['network']=net

# ---- D. remodeling index ----
S['STRUCT_index']=S[['ECM_CROSS','HIF_MATRIX','TGFB','YAPTAZ']].mean(1)
S['INFLAM_index']=S[['MACRO','MONO_RECRUIT','HYPOXIA']].mean(1)
S['REMODEL_index']=S['STRUCT_index']-S['INFLAM_index']
u,p=stats.mannwhitneyu(S.loc[SAT,'REMODEL_index'],S.loc[VAT,'REMODEL_index'])
results['remodeling_index']={'SAT_mean':round(float(S.loc[SAT,'REMODEL_index'].mean()),3),
    'VAT_mean':round(float(S.loc[VAT,'REMODEL_index'].mean()),3),'p':float(p)}

S.to_csv(OUT+"GTEx_module_scores_per_sample.csv")
with open(OUT+"GTEx_replication_results.json","w") as f: json.dump(results,f,indent=1,default=str)

# ---- pretty print ----
print("\n===== A. DEPOT CONTRAST (SAT subcut vs VAT omentum) =====")
print(f"{'module':14s} {'SAT':>7s} {'VAT':>7s} {'dir':>8s} {'rb':>7s} {'p':>10s}")
for r in contrast:
    print(f"{r['module']:14s} {r['SAT_mean']:7.3f} {r['VAT_mean']:7.3f} {r['dir']:>8s} {r['rank_biserial']:7.3f} {r['p']:.2e}")
print("\n===== B. REGULATOR ~ LOX/POSTN (within depot, Spearman) =====")
for r in reg_rows:
    print(f"{r['depot']:3s} {r['regulon']:12s} ~ {r['outcome']:10s} rho={r['rho']} p={r['p']:.2e} n={r['n']}")
print("\n===== C. NETWORK =====")
for d in ['SAT','VAT']:
    print(f"-- {d} --")
    for k,v in net[d].items(): print(f"   {k:22s} rho={v[0]} p={v[1]:.2e} n={v[2]}")
print("\n===== D. REMODELING INDEX =====")
print(results['remodeling_index'])
print("\nDONE")
```
