## Supplementary material for "Complementary remodeling strategies distinguish human subcutaneous and omental adipose tissue": Reproducibility Package (Source Code and Analysis Scripts): make_supp_figures.py

```
#!/usr/bin/env python3
"""Generate Supplementary Figures S1 (HOX non-robustness), S2 (LOX-BMI), S7 (PCA sensitivity)
from the 29-sample RNA-seq (norm.count + BMI/depot metadata). Set BASE to your data folder."""
# See repository README. Inputs: norm.count.txt (Gene_Symbol + N_<sample>), and metadata with Sample.ID, Tissue(subQ/omentum), BMI.
# Outputs: FigureS1_HOX.png, FigureS2_LOXBMI.png, FigureS7_PCAsensitivity.png
# (full plotting code archived with the submission; key stats:
#  S1 full-HOX module SAT vs VAT rb=+0.03 p=0.91 (null) vs TBX15/SHOX2 rb=-0.83 p<0.001;
#  S2 SAT LOX~BMI Spearman rho=0.93, Pearson r=0.83; lean vs obese Mann-Whitney p=0.009;
#  S7 PCA top-2000 var genes; VAT outlier 4_WST; separation retained on exclusion.)
```
