## Supplementary material for "Complementary remodeling strategies distinguish human subcutaneous and omental adipose tissue": Reproducibility Package (Source Code and Analysis Scripts): mechanism_analysis.py

```
#!/usr/bin/env python3
import pandas as pd, numpy as np, json, warnings
from scipy import stats
warnings.filterwarnings('ignore'); np.random.seed(1)
import argparse, os
_ap = argparse.ArgumentParser(description="n=29 module-score / mechanism analysis")
_ap.add_argument("--cohort-dir", default=os.environ.get("COHORT_DIR", "./cohort"),
                 help="directory with norm.count.txt and the CT-RNA xlsx (deposited on Zenodo/GEO)")
_ap.add_argument("--out", default=os.environ.get("OUT_DIR", "./outputs"))
_args, _ = _ap.parse_known_args()
BASE = _args.cohort_dir.rstrip("/") + "/"
OUT = _args.out.rstrip("/") + "/"
os.makedirs(OUT, exist_ok=True)
COUNT=BASE+"norm.count.txt"; XLSX=BASE+"RNA20_corrected_CT_RNA_reanalysis_20260626.xlsx"
c=pd.read_csv(COUNT,sep='\t'); c.columns=[x.replace('N_','',1) for x in c.columns]
c=c.dropna(subset=['Gene_Symbol']); sc=[x for x in c.columns if x not in ('Gene_ID','Gene_Symbol')]
c['m']=c[sc].mean(1); c=c.sort_values('m',ascending=False).drop_duplicates('Gene_Symbol')
log2=np.log2(c.set_index('Gene_Symbol')[sc]+1)
import openpyxl
wb=openpyxl.load_workbook(XLSX,data_only=True); r=list(wb['RNA_CT_integrated'].iter_rows(values_only=True))
hdr=list(r[2]); meta=pd.DataFrame([row for row in r[3:] if row[0] is not None],columns=hdr).set_index('Sample.ID')
for col in ['Age','BMI','new_VAT_area_cm2','new_SAT_area_cm2','new_VAT_SAT_ratio','new_VAT_HU_Mean','new_SAT_HU_Mean']:
    meta[col]=pd.to_numeric(meta[col],errors='coerce')
meta['depot']=meta['Tissue'].map({'subQ':'SAT','omentum':'VAT'})
common=[s for s in log2.columns if s in meta.index]; log2=log2[common]; meta=meta.loc[common]
SAT=meta.index[meta.depot=='SAT'].tolist(); VAT=meta.index[meta.depot=='VAT'].tolist()
print(f"samples: SAT={len(SAT)} VAT={len(VAT)} total={len(common)}")
Z=log2.sub(log2.mean(1),axis=0).div(log2.std(1,ddof=1).replace(0,np.nan),axis=0)
def modz(genes,cols):
    gg=[g for g in genes if g in Z.index]; return Z.loc[gg,cols].mean(0),gg
HYPOXIA=['VEGFA','SLC2A1','LDHA','PGK1','ADM','PDK1','BNIP3','ANGPTL4','EGLN3','NDRG1','ALDOA','ENO1','HK2','PFKFB3']
HIF_MATRIX=['P4HA1','P4HA2','PLOD1','PLOD2','LOX','LOXL2']
TGFB=['SERPINE1','SMAD7','TGFBI','SKIL','JUNB','LTBP2','TGFB1','TGFBR1','THBS1','CCN2']
YAPTAZ=['CCN2','CCN1','AMOTL2','AXL','THBS1','F3','TAGLN','ACTA2','MYL9','TEAD1']
ECM_CROSS=['LOX','LOXL1','LOXL2','POSTN','P4HA1','PLOD2']
COLLAGEN=['COL1A1','COL1A2','COL3A1','COL6A3']
MACRO=['CD68','CD163','MRC1','MSR1','CSF1R','ITGAM','AIF1','TYROBP','C1QA','C1QB']
MONO_RECRUIT=['CCL2','CCR2','CCL7','CCL8','CX3CR1','S100A8','S100A9']
ANGIO=['VEGFA','KDR','FLT1','PECAM1','NRP2','APLNR','CCN1','ESM1']
DEVEL=['TBX15','SHOX2','HOXA5','HOXA10','NR2F1']; MESO=['UPK3B','KRT19']
modules={'HYPOXIA':HYPOXIA,'HIF_MATRIX':HIF_MATRIX,'TGFB':TGFB,'YAPTAZ':YAPTAZ,'ECM_CROSS':ECM_CROSS,
 'COLLAGEN':COLLAGEN,'MACRO':MACRO,'MONO_RECRUIT':MONO_RECRUIT,'ANGIO':ANGIO,'DEVEL':DEVEL,'MESO':MESO}
S={}; used={}
for k,gs in modules.items(): S[k],used[k]=modz(gs,common)
S=pd.DataFrame(S,index=common); S['depot']=meta['depot']; S['BMI']=meta['BMI']
for col in ['new_VAT_area_cm2','new_SAT_area_cm2','new_VAT_SAT_ratio','new_VAT_HU_Mean','new_SAT_HU_Mean','Age']: S[col]=meta[col]
for gname in ['LOX','POSTN','LOXL2','COL1A1','COL3A1','COL6A3','HIF1A','EPAS1','TGFB1','WWTR1','YAP1','PIEZO1','PTK2','CCN2','CD68','CD163']:
    if gname in log2.index: S[gname+'_expr']=log2.loc[gname,common]
def spear(a,b):
    m=(~pd.isna(a))&(~pd.isna(b)); n=int(m.sum())
    if n<4: return (np.nan,np.nan,n)
    rho,p=stats.spearmanr(a[m],b[m]); return (round(float(rho),3),round(float(p),4),n)
results={'genes_used':used}
reg_rows=[]
for depot,cols in [('SAT',SAT),('VAT',VAT)]:
    sub=S.loc[cols]
    for reg in ['HYPOXIA','TGFB','YAPTAZ','HIF_MATRIX']:
        for outc in ['LOX_expr','POSTN_expr']:
            rho,p,n=spear(sub[reg],sub[outc]); reg_rows.append(dict(depot=depot,regulon=reg,outcome=outc,rho=rho,p=p,n=n))
    for tf in ['HIF1A_expr','EPAS1_expr','TGFB1_expr','WWTR1_expr','YAP1_expr','PIEZO1_expr','PTK2_expr']:
        if tf in sub: rho,p,n=spear(sub[tf],sub['LOX_expr']); reg_rows.append(dict(depot=depot,regulon=tf,outcome='LOX_expr',rho=rho,p=p,n=n))
results['upstream_regulator']=reg_rows
def partial_spear(x,y,z):
    df=pd.concat([x,y,z],axis=1).dropna(); df.columns=['x','y','z']
    if len(df)<5: return (np.nan,np.nan,len(df))
    rx=stats.rankdata(df['x']); ry=stats.rankdata(df['y']); rz=stats.rankdata(df['z'])
    def resid(a,b):
        b1=np.vstack([np.ones_like(b),b]).T; beta=np.linalg.lstsq(b1,a,rcond=None)[0]; return a-b1@beta
    ex=resid(rx,rz); ey=resid(ry,rz); rho,p=stats.pearsonr(ex,ey); return (round(float(rho),3),round(float(p),4),len(df))
med={}; sat=S.loc[SAT]
for outc in ['LOX_expr','POSTN_expr','ECM_CROSS']:
    med[outc]=dict(BMI_to_out=spear(sat['BMI'],sat[outc]),BMI_to_hypoxia=spear(sat['BMI'],sat['HYPOXIA']),
        hypoxia_to_out=spear(sat['HYPOXIA'],sat[outc]),BMI_to_out_ctrl_hypoxia=partial_spear(sat['BMI'],sat[outc],sat['HYPOXIA']))
results['mediation_SAT']=med
ov=[]
for outc in ['new_VAT_SAT_ratio','new_VAT_area_cm2']:
    for pred in ['ECM_CROSS','LOX_expr','POSTN_expr']:
        rho,p,n=spear(sat[pred],sat[outc]); ov.append(dict(predictor_in_SAT=pred,outcome=outc,rho=rho,p=p,n=n))
results['overflow_from_SAT']=ov
net={}
for depot,cols in [('SAT',SAT),('VAT',VAT)]:
    sub=S.loc[cols]
    net[depot+'_key']={'hypoxia~ECMcross':spear(sub['HYPOXIA'],sub['ECM_CROSS']),
        'hypoxia~macro':spear(sub['HYPOXIA'],sub['MACRO']),'ECMcross~macro':spear(sub['ECM_CROSS'],sub['MACRO']),
        'macro~mono_recruit':spear(sub['MACRO'],sub['MONO_RECRUIT']),'ECMcross~TGFB':spear(sub['ECM_CROSS'],sub['TGFB']),
        'ECMcross~YAPTAZ':spear(sub['ECM_CROSS'],sub['YAPTAZ'])}
results['network']=net
contrast=[]
for k in ['HYPOXIA','HIF_MATRIX','ECM_CROSS','COLLAGEN','MACRO','MONO_RECRUIT','TGFB','YAPTAZ','ANGIO','DEVEL','MESO','LOX_expr','POSTN_expr']:
    a=S.loc[SAT,k].dropna(); b=S.loc[VAT,k].dropna(); u,p=stats.mannwhitneyu(a,b,alternative='two-sided')
    rb=1-2*u/(len(a)*len(b))
    contrast.append(dict(module=k,SAT_mean=round(float(a.mean()),3),VAT_mean=round(float(b.mean()),3),
        dir='SAT>VAT' if a.mean()>b.mean() else 'VAT>SAT',rank_biserial=round(float(rb),3),p=round(float(p),4)))
results['depot_contrast']=contrast
S['STRUCT_index']=S[['ECM_CROSS','HIF_MATRIX','TGFB','YAPTAZ']].mean(1)
S['INFLAM_index']=S[['MACRO','MONO_RECRUIT','HYPOXIA']].mean(1)
S['REMODEL_index']=S['STRUCT_index']-S['INFLAM_index']
dri=[]
for depot,cols in [('SAT',SAT),('VAT',VAT)]:
    dri.append(dict(depot=depot,STRUCT=round(float(S.loc[cols,'STRUCT_index'].mean()),3),
        INFLAM=round(float(S.loc[cols,'INFLAM_index'].mean()),3),REMODEL=round(float(S.loc[cols,'REMODEL_index'].mean()),3)))
u,p=stats.mannwhitneyu(S.loc[SAT,'REMODEL_index'],S.loc[VAT,'REMODEL_index'])
results['remodeling_index']={'by_depot':dri,'SAT_vs_VAT_p':round(float(p),6)}
for depot,cols in [('SAT',SAT),('VAT',VAT)]:
    results['remodeling_index'][f'{depot}_STRUCT~BMI']=spear(S.loc[cols,'STRUCT_index'],S.loc[cols,'BMI'])
    results['remodeling_index'][f'{depot}_INFLAM~BMI']=spear(S.loc[cols,'INFLAM_index'],S.loc[cols,'BMI'])
S.to_csv(OUT+"module_scores_per_sample.csv")
with open(OUT+"mechanism_results.json","w") as f: json.dump(results,f,indent=1,default=str)
print("DONE")
```
