## Supplementary figures and images for "Complementary remodeling strategies distinguish human subcutaneous and omental adipose tissue"

### Figure1.png

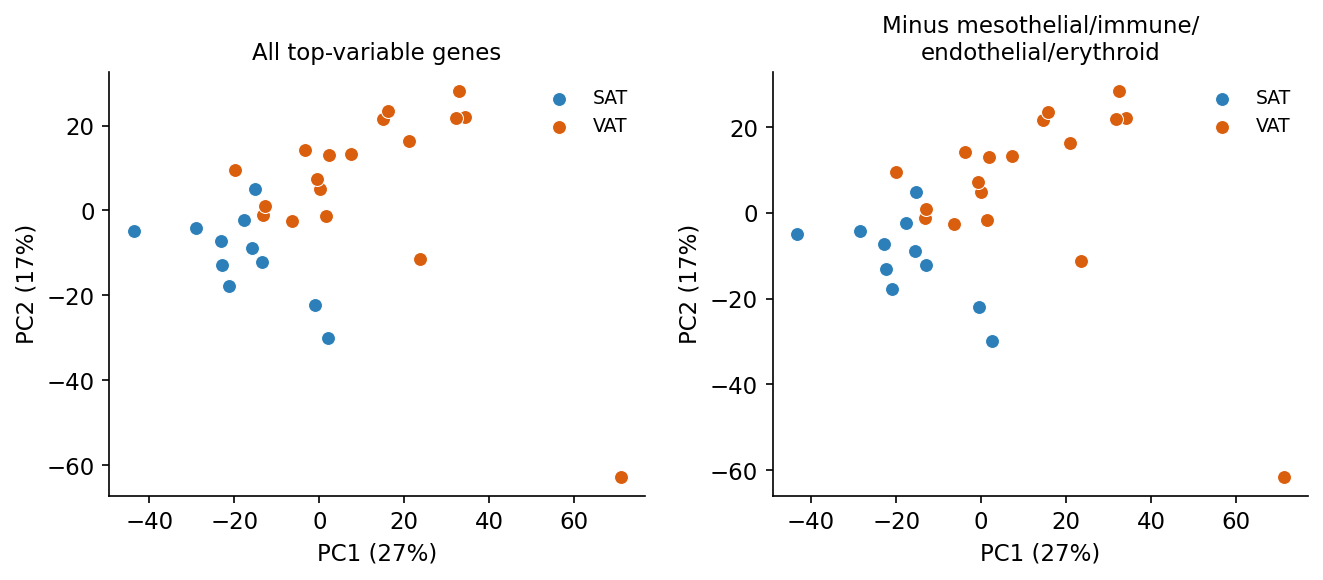

### Figure2.png

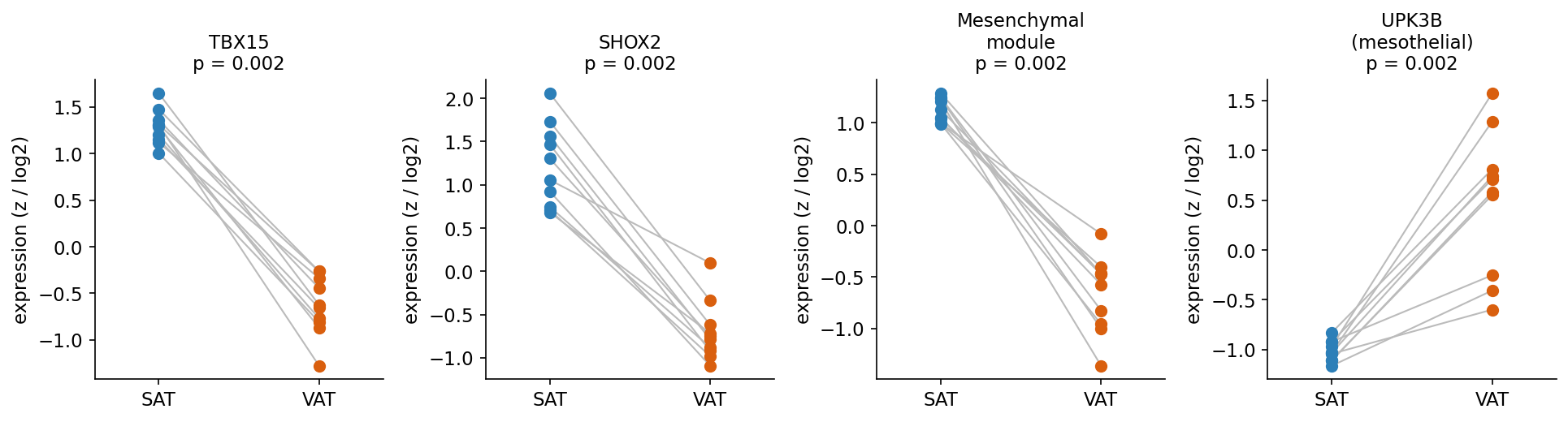

### Figure3_histology.png

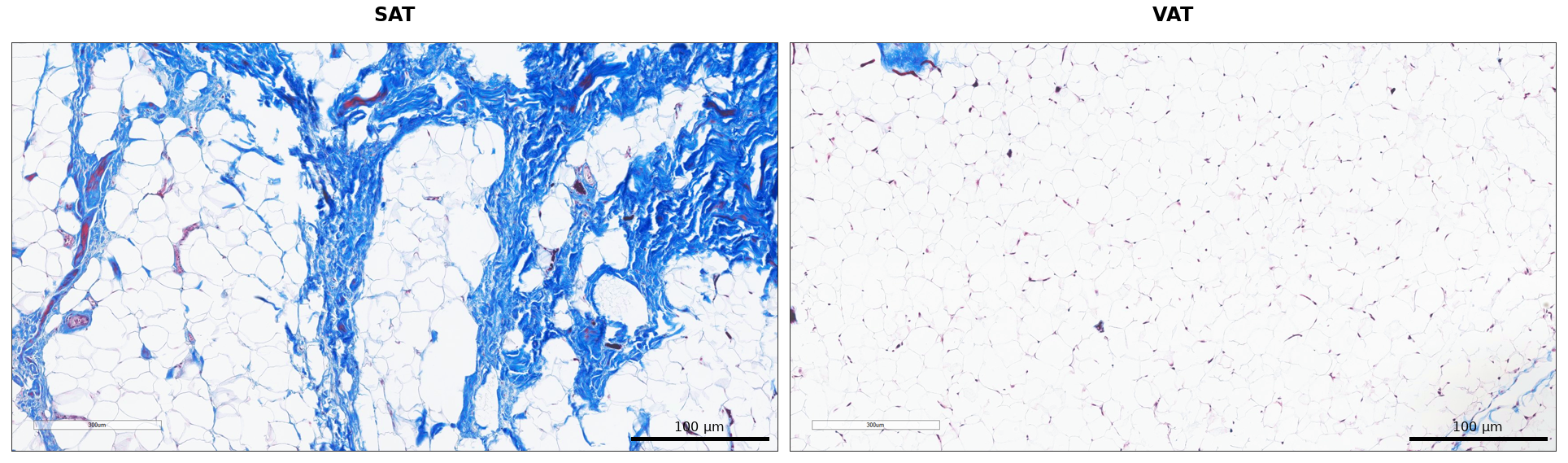

### Figure3_panelB.png

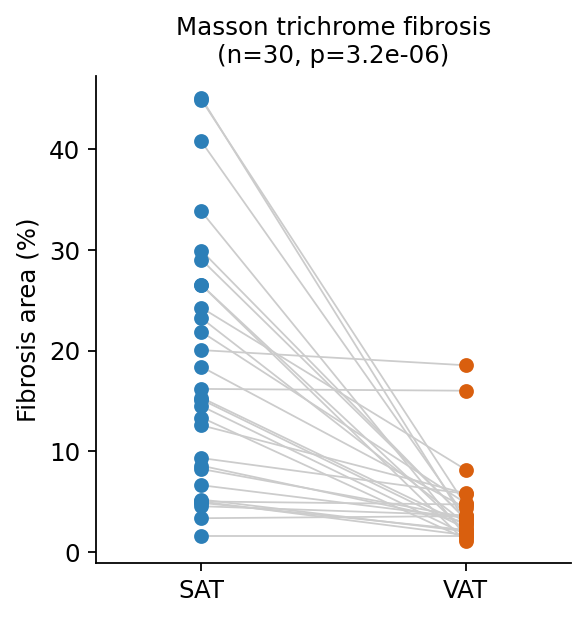

### Figure4_model.png

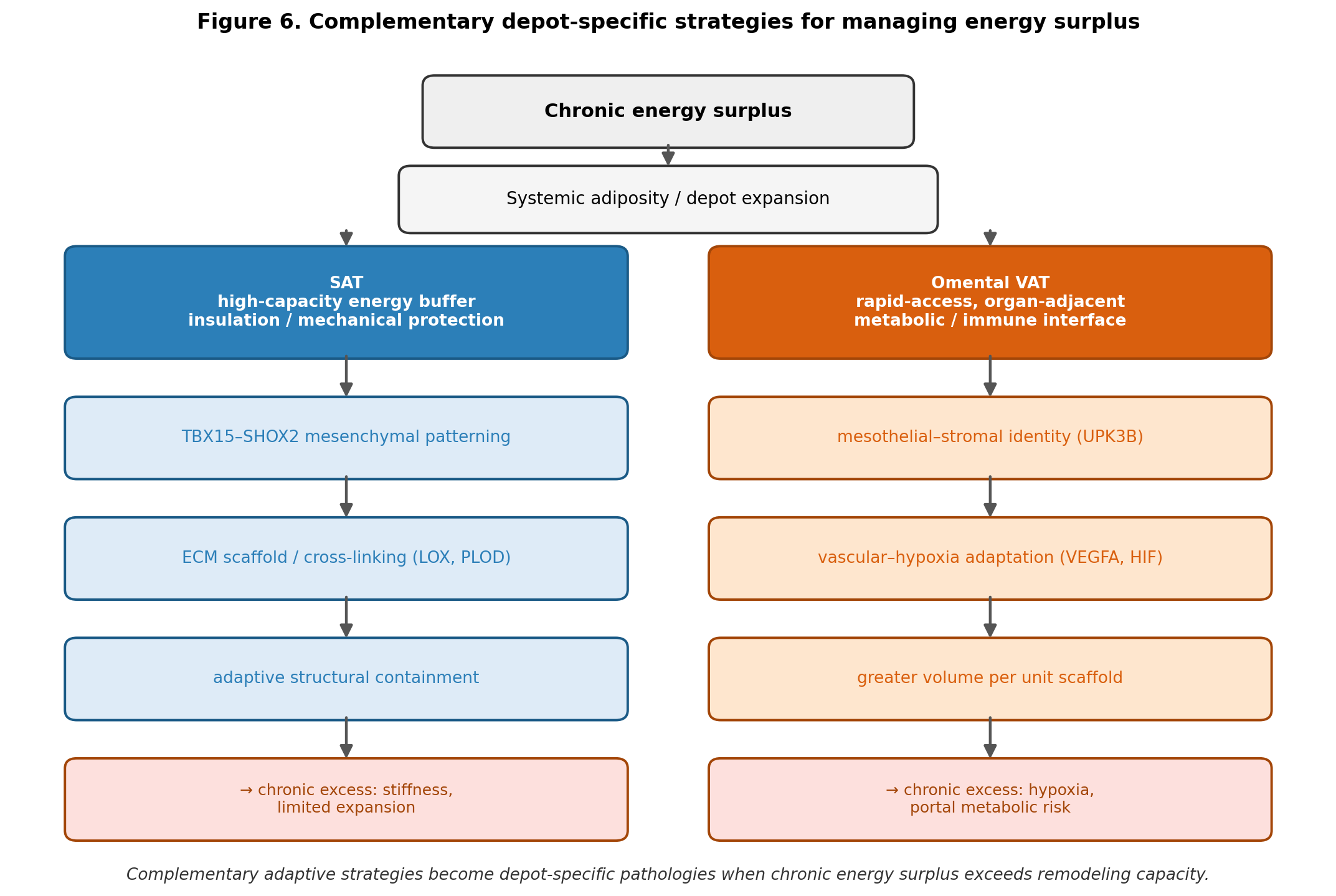

### Figure5_storage.png

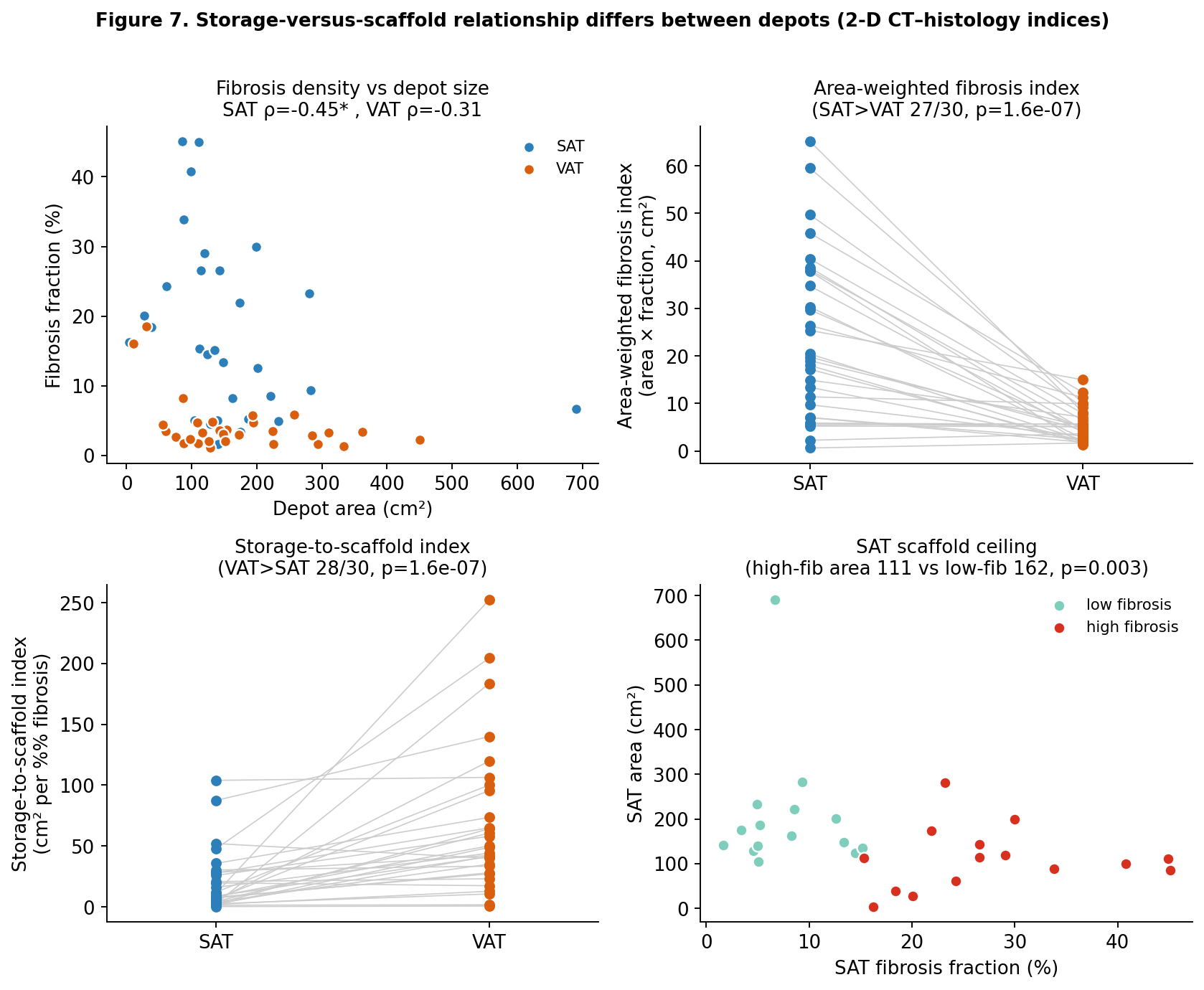

### Figure6_external_validation.png

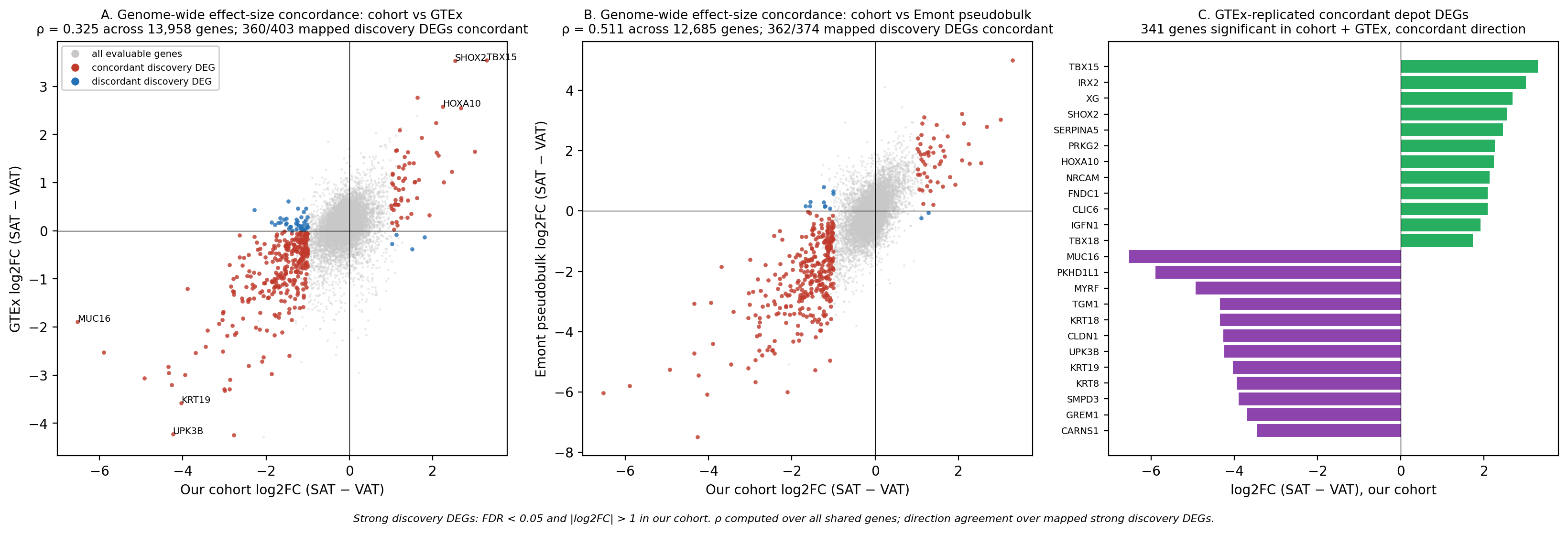

### FigureS1_HOX.png

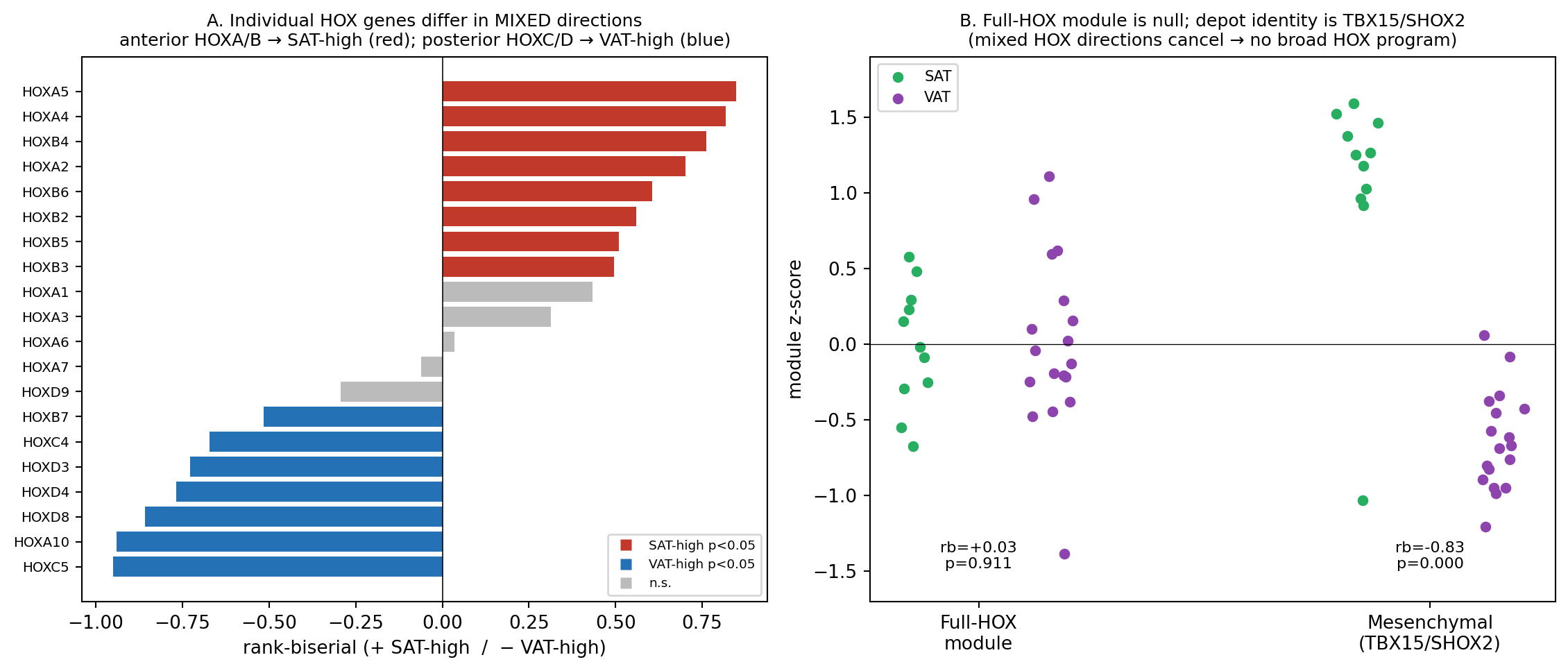

### FigureS2_LOXBMI.png

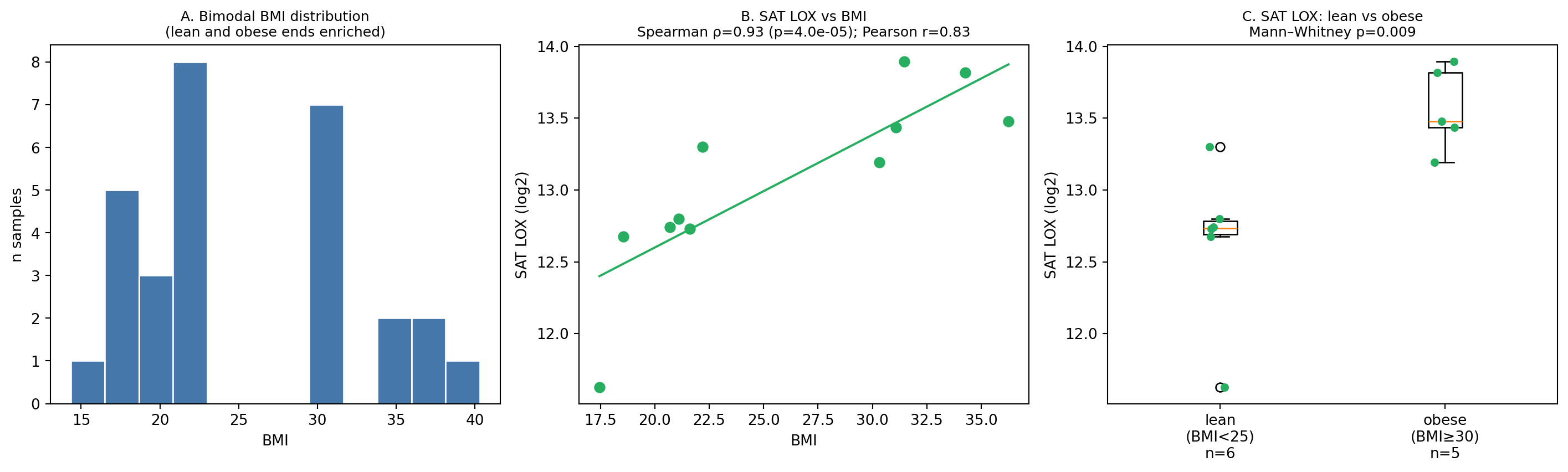

### FigureS3_LOOclassifier.png

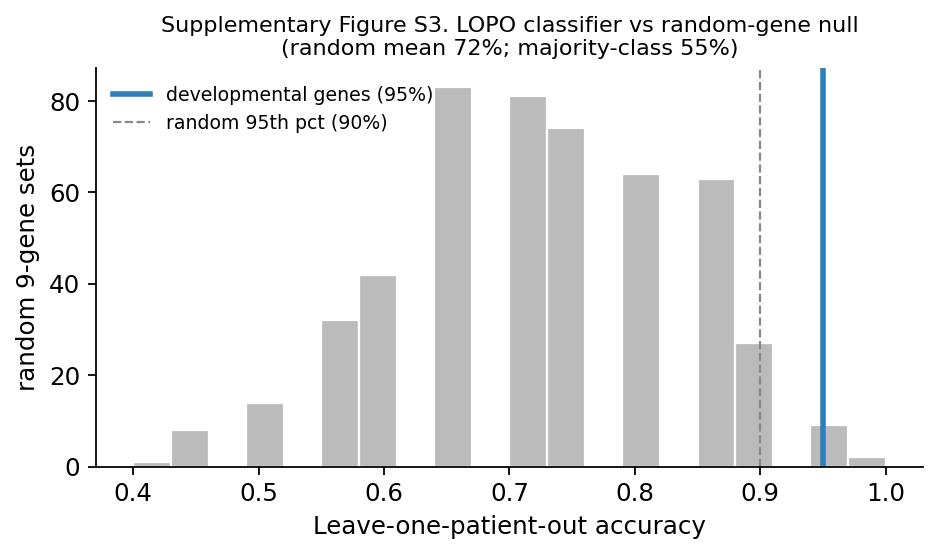

### FigureS4_adjustment.png

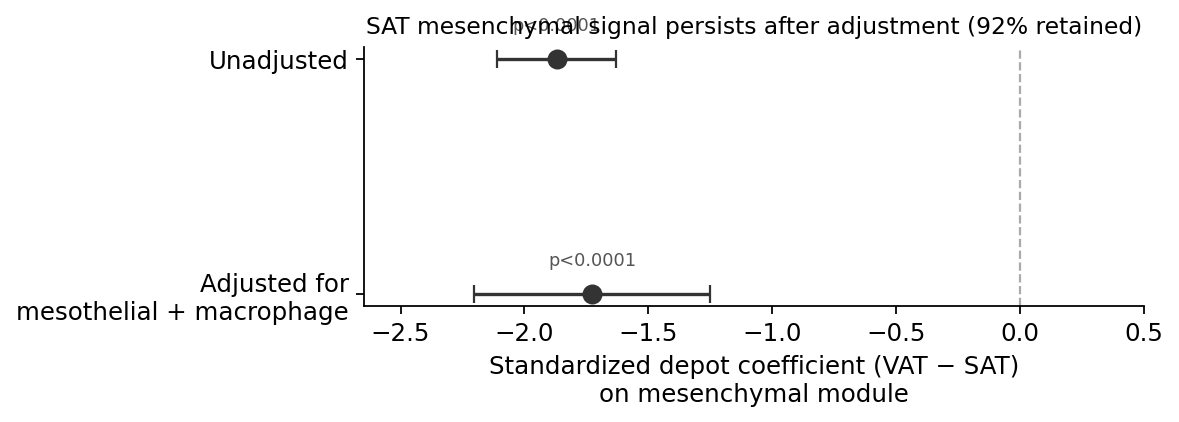

### FigureS5_CD68.png

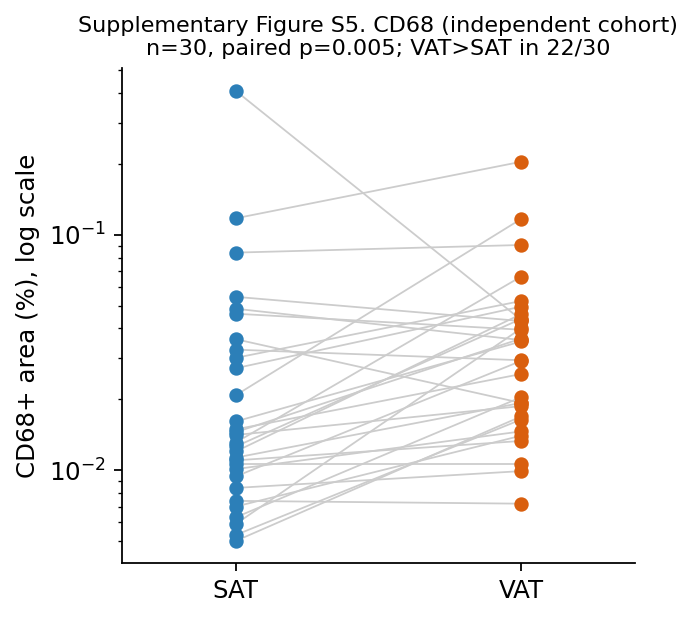

### FigureS6_CTRNA.png

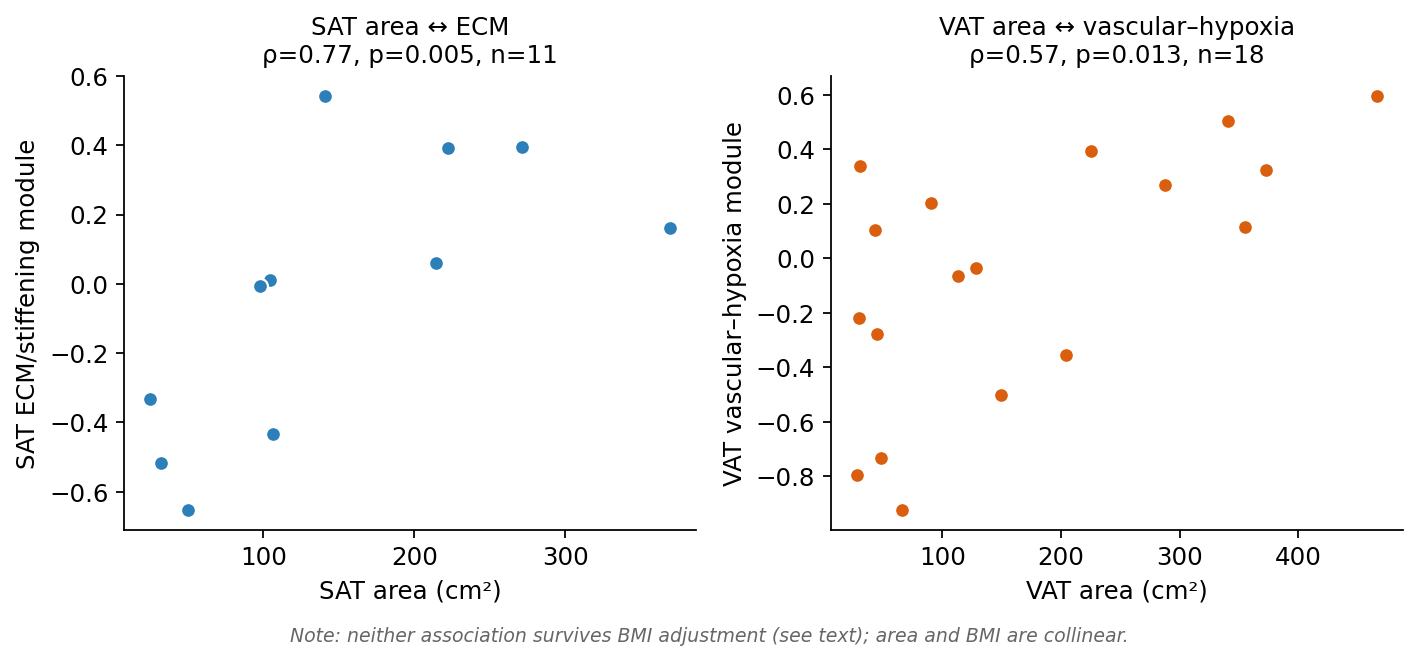

### FigureS7_PCAsensitivity.png

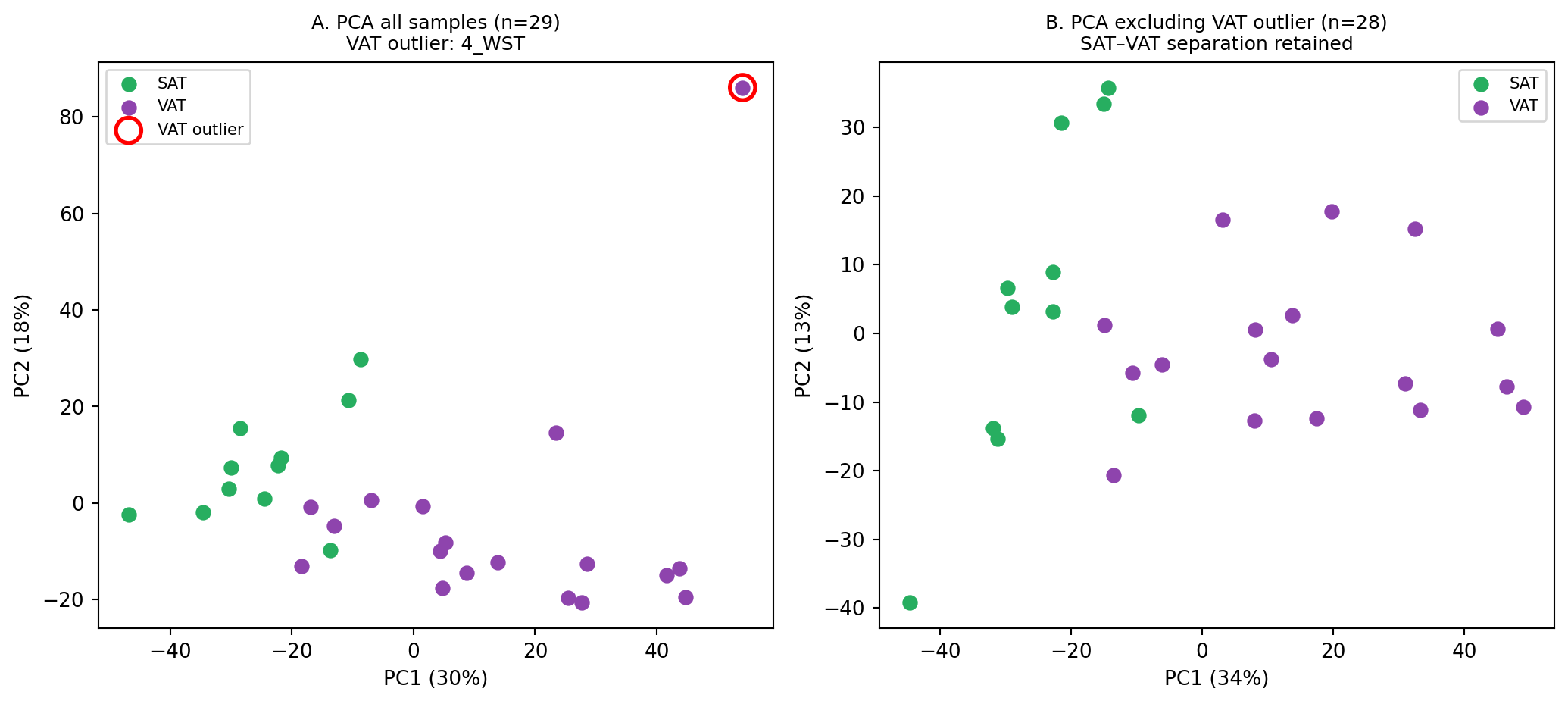

### FigureS8_HU.png

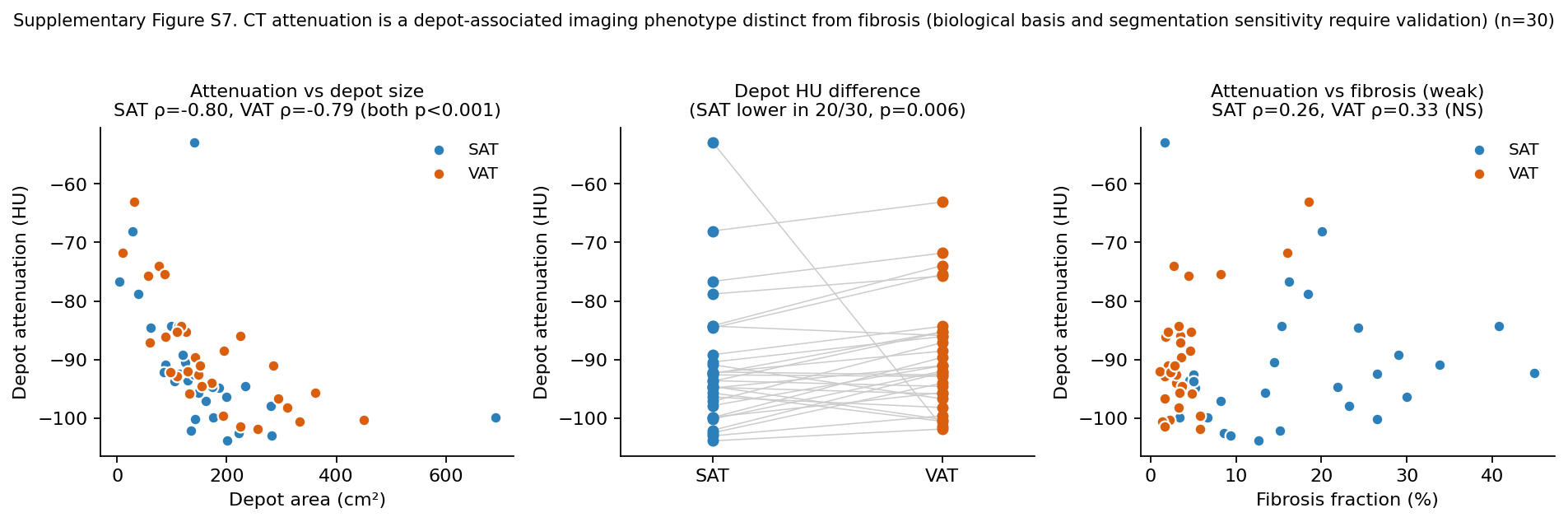

### FigureS9_cellorigin.png

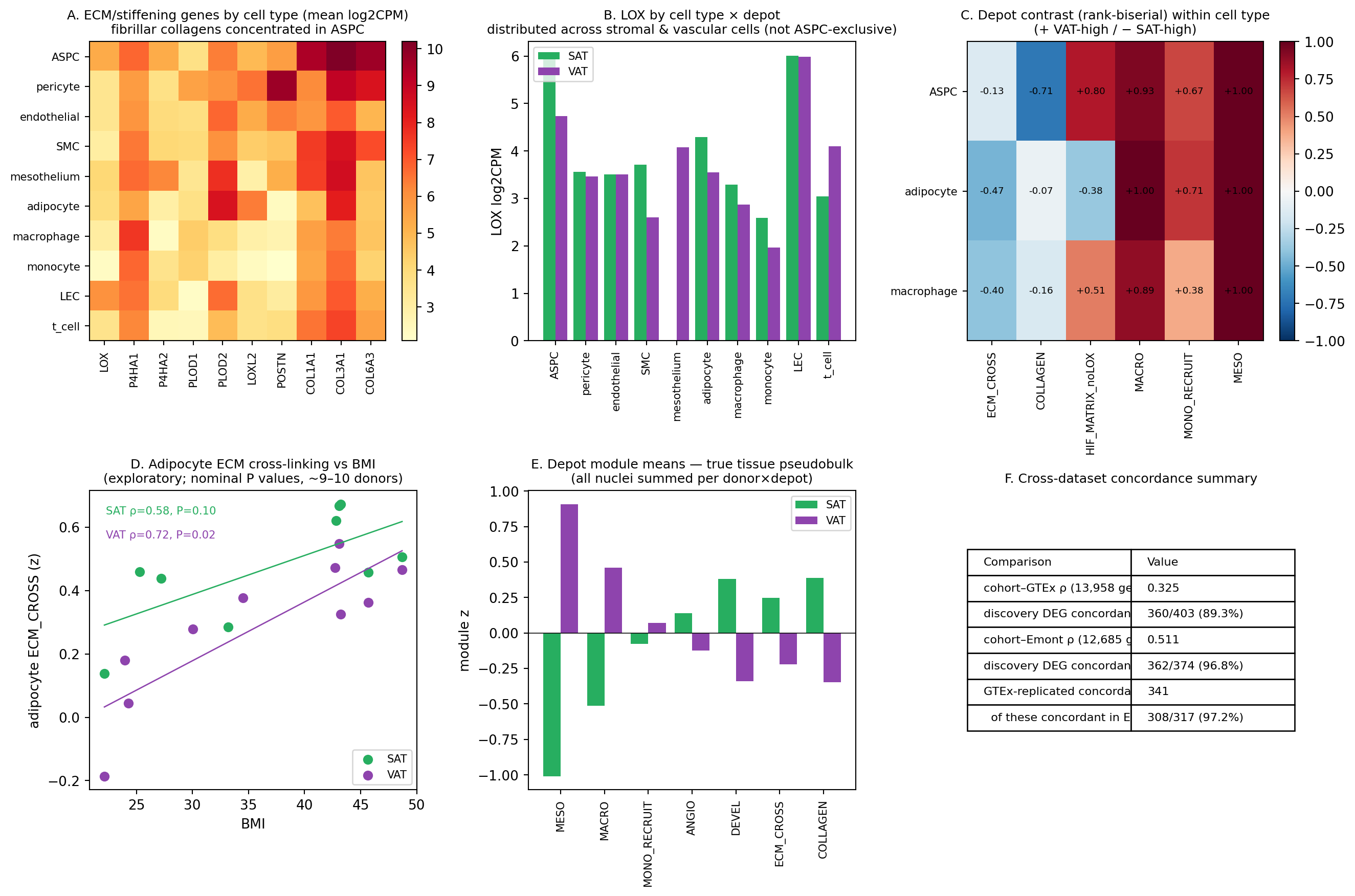
