## Supplementary Information for "Complementary remodeling strategies distinguish human subcutaneous and omental adipose tissue"

### Supplementary Methods

#### S-Methods 1. RNA sequencing and processing (full detail)

Total RNA was extracted from adipose tissue and sequenced by a commercial provider (Macrogen, Seoul, Republic of Korea). RNA was quantified with the Quant-iT RiboGreen assay (Invitrogen) and integrity/size distribution assessed on an Agilent 2100 Bioanalyzer/4200 TapeStation. Because adipose specimens yielded low-quality, partially degraded RNA, sample eligibility was assessed by DV200 rather than RIN, and a number of specimens required re-extraction and re-submission before passing quality control. Sequencing libraries were prepared with the Agilent SureSelectXT RNA Direct Library Preparation Kit—a capture-based protocol suited to lower-integrity input—and passed library QC by qPCR and TapeStation D1000. Libraries were sequenced on an Illumina platform as paired-end 151-bp reads (2 × 151 bp) for 29 samples. Reads were processed against the human reference genome GRCh38 and quantified against RefSeq gene annotation, yielding raw counts for 46,427 genes. Genes with a zero count in at least one of the 29 samples were excluded, leaving 16,493 genes for statistical analysis. Differential expression was performed with the edgeR R package: library sizes were normalized by the trimmed mean of M-values (TMM) method (calcNormFactors), and the depot contrast (all 29 samples, treated as two groups) was tested with edgeR's exactTest and reported at $|\text{fold change}| \geq 2$ with unadjusted $p < 0.05$ as an exploratory discovery threshold; the complementary within-patient paired comparison (10 pairs) is reported separately (Supplementary Table S2), and the external-validation discovery set was defined by FDR < 0.05 and $|\log2FC| > 1$ (Section 2.9). For visualization, log2(CPM + 1) and TMM-normalized values were used. For the depot-comparison analyses reported here, expression was quantified for 16,493 gene symbols across the 29-sample set; for duplicated gene symbols, the row with the highest mean expression was retained. Sequencing was performed across more than one submission batch (Supplementary Table S4).

#### S-Methods 2. Module definitions

Gene modules were defined a priori (Supplementary Table S1). Each module score was the mean, across the genes in the module, of cohort-wide per-gene z-scores (standardized across all 29 samples). Identity modules: mesenchymal patterning (TBX15, TBX18, SHOX2, MEOX1, SATB2, IRX2), mesothelial (UPK3B, KRT19, KRT8, KRT18, MUC16, CALB2), and—for completeness only—HOX positional (HOXA5, HOXA10, HOXD4). Remodeling modules: ECM stiffening (LOX, LOXL1/2, POSTN, PLOD1/2, P4HA1/2), vascular–hypoxia (HIF1A, EPAS1, VEGFA, SLC2A1, PGK1, LDHA, CA9, PECAM1, CDH5, VWF), and macrophage (CD68,

CD163, MRC1, MSR1). Genes not detected in the matrix were omitted; module composition as analyzed is listed in Supplementary Table S1.

#### S-Methods 3. Statistical analysis (full detail)

Paired SAT–VAT comparisons used the Wilcoxon signed-rank test (exact) on within-patient differences (primary), with matched-pairs rank-biserial effect sizes; group comparisons used the Mann–Whitney U test with rank-biserial effect sizes. Identity signals were re-tested at the single-gene level and with alternative gene selections, including the full complement of detected HOX genes, to assess robustness. Linear mixed-effects models were fitted on the 20 specimens from the 10 paired patients (module ~ depot + covariates, with a patient random intercept); models were fitted by maximum likelihood, and coefficients are reported as standardized β with 95% CI, standard error, exact p, and random-effect variance in Supplementary Table S3. Cell-composition sensitivity was evaluated by adding mesothelial and macrophage module scores as fixed-effect covariates (module ~ depot + mesothelial + macrophage + (1|patient)); no interaction terms were included. Adiposity confounding was evaluated by adding BMI and age (module ~ depot + BMI + age + (1|patient)). Retained effect was defined as $100 \times |\beta_{adjusted}| / |\beta_{unadjusted}|$. Principal component analysis was performed on the top 2,000 most variable genes (by variance) after standardization, as a descriptive/sensitivity analysis: (i) on all top-variable genes, (ii) after excluding mesothelial/immune/endothelial/erythroid genes (list in Supplementary Table S1) to reduce the contribution of major cell-composition-associated transcripts, and (iii) on identity genes alone. Because PC sign is arbitrary, depot separation is reported as the absolute rank-biserial effect size of the best-separating component, and the accompanying p-values are descriptive. A leave-one-patient-out classifier (Supplementary Figure S3) was evaluated against a random-gene null distribution. CT–RNA associations used Spearman correlation and BMI-adjusted partial correlation and multivariable regression. Missing clinical values were handled by pairwise deletion (each analysis used all samples with complete data for the variables involved; per-variable availability is given in Supplementary Table S1). Analyses used Python (pandas, numpy, scipy, statsmodels, scikit-learn); a reproduction package regenerating every reported statistic is provided (Section: Data availability).

#### S-Methods 4. Primary integrative analyses and CT–histology indices

A small set of directional hypotheses was designated as primary (these were specified analytically before integration but were not registered in a formal protocol, and are therefore described as primary rather than pre-registered): SAT area ↔ SAT ECM module; VAT area ↔ VAT vascular–hypoxia module; and, at the tissue level, whether the independent histologic cohort showed depot fibrosis patterns directionally concordant with the transcriptomic ECM program. Because only one patient overlapped between the RNA and histology cohorts, no patient-level RNA–histology association was tested; histology provides independent, group-level support only. All other gene–CT and metabolic associations were exploratory. For the histologic cohort, two 2-D, area-weighted CT–histology surrogates were defined per depot from the single-slice CT area (A, $cm^2$) and the histologic fibrosis area fraction (f, %). The area-weighted fibrosis index = $A \times (f/100)$, with units of $cm^2$ (an estimate of fibrotic cross-sectional area on the analyzed slice). The storage-to-scaffold index = A / f, with units of $cm^2$ per percentage-point of fibrosis (higher values denote more adipose area per unit fractional fibrosis). Both are 2-D surrogates computed from a single CT slice and a separate histologic section; they are not measurements of tissue volume, storage capacity, or collagen mass, and are labelled as indices throughout. The between-depot ratio of the storage-to-scaffold index is invariant to whether f is expressed as a percentage or a fraction.

#### S-Methods 5. Histology (staining, imaging, classifier)

Histologic analyses were performed in an independent cohort (n = 30 with paired specimens) that was essentially non-overlapping with the RNA-seq cohort (overlap = 1 patient). Formalin-fixed, paraffin-embedded specimens were sectioned at 4 µm. Masson trichrome staining was performed on an automated platform (BenchMark Special Stains, Ventana Medical Systems, Tucson, AZ, USA), staining collagen blue, nuclei black, and cytoplasm/muscle/keratin red. CD68 immunohistochemistry was performed on a BenchMark ULTRA PLUS system with a rabbit monoclonal anti-CD68 antibody (clone SP251, Ventana/Roche), heat-induced antigen retrieval (CC1, 32 min), antibody incubation (16 min), and OptiView DAB detection. Whole-slide images were analyzed in QuPath v0.5.1; the adipose region was manually delineated after excluding mesothelial surfaces, hemorrhage, large vessels, background, and artifacts. Fibrosis was quantified using a supervised pixel classifier generated from representative fibrotic and non-fibrotic regions in a representative subcutaneous adipose tissue whole-slide image. The identical classifier was subsequently applied to all SAT and VAT specimens to calculate the fibrosis area fraction (%). Because a classifier trained on a single subcutaneous slide could in principle be less sensitive to the thinner, more lightly stained collagen bundles of omental VAT, we regard the depot fibrosis comparison as robust in direction but potentially conservative in the VAT estimate; we note, however, that measured VAT fibrosis was not floored (range 1.1–18.6%, with 5/30 specimens > 5%), arguing against complete failure to detect VAT collagen, although proportional underestimation cannot be excluded. A classifier trained on both depots and pathologist-annotated concordance are planned to confirm the magnitude (Limitations). CD68 was quantified with a threshold-based pixel classifier applied uniformly (CD68-positive area fraction, %). Representative images shown are from a single patient with fibrosis values near the cohort median; a full low/median/high representative panel will be assembled from the digitized slide archive for the revised submission. Images were acquired at 100× total magnification (10× objective); scale bars annotated on the images denote 300 µm.

#### S-Methods 6. CT analysis and attenuation review

Preoperative non-contrast abdominal computed tomography (CT) scans obtained within 30 days before surgery were analyzed for measurement of SAT and VAT. A single axial CT image at the level of the inferior end plate of the third lumbar vertebra (L3) was selected for all patients, as this anatomical landmark is widely used for quantitative assessment of abdominal adipose tissue. CT images were analyzed using the open-source software platform 3D Slicer (version 5.10.0; Slicer community). VAT and SAT were manually segmented on the selected L3 slice using a predefined attenuation threshold of -190 to -30 Hounsfield units (HU). Manual correction was performed when necessary to exclude non-adipose tissue structures and ensure accurate tissue delineation. The cross-sectional areas of VAT and SAT were calculated automatically using the Segment Cross-Section Area module and expressed in square centimeters (cm2). Mean and median attenuation values (HU) for VAT and SAT were obtained using the Segment Statistic module. Attenuation values were reviewed for physiologic plausibility. In the histologic cohort (n = 30), CT area and HU were complete for all patients and all values fell within the expected adipose range (SAT and VAT HU −53 to −104); in the separate RNA cohort, a small number of specimens showed atypically high mean HU inconsistent with pure adipose tissue (e.g., one subcutaneous measurement at −14.8 HU), likely reflecting partial-volume inclusion of fascia/muscle or a segmentation artifact (flagged in Supplementary Table S1b, and undergoing source-image recheck). HU-based analyses are reported as exploratory. Post-operative weight change, follow-up CT, and metabolic outcomes were not analyzed and are reserved for a separate longitudinal study.

#### S-Methods 7. External-validation pipeline

To assess generalizability, depot-specific transcriptional programs were compared with two external human datasets. For external comparison, discovery depot DEGs were defined from the 29-sample normalized counts (Welch's t-test; FDR < 0.05 and |log2FC| > 1; 420 genes), a set distinct from the primary edgeR contrast (Section 2.3). GTEx v8 gene-level TPM for adipose–subcutaneous (n = 663) and adipose–visceral/omentum (n = 541) [25] were log2(TPM+1)-transformed and tested with Welch's t-test and Benjamini–Hochberg FDR. The human single-nucleus atlas of Emont et al. [12] (137,684 nuclei, 29,093 genes; Single Cell Portal SCP1376) was aggregated by sparse matrix multiplication to 273 donor × depot × cell-type pseudobulks (the 230 with ≥ 10 nuclei were used for module-level analyses); counts were CPM-normalized and log2-transformed. Genome-wide depot fold-changes in the atlas were computed on paired donors (n = 6; Wilcoxon signed-rank), and module z-scores were compared within cell types (Mann–Whitney, rank-biserial). Cross-dataset concordance was quantified as the Spearman correlation of depot log2 fold-changes over shared genes and the proportion of discovery DEGs evaluable in each dataset with matching direction. GTEx adipose–visceral(omentum) corresponds to the omental VAT sampled here. Analysis scripts and processed matrices are provided (Data availability).

### Supplementary Results

#### S-Results 1. HOX positional program is not a robust depot-identity axis

By contrast, a broad HOX program did not robustly distinguish the depots. Although a three-gene HOX module (HOXA5/HOXA10/HOXD4) reached significance (paired p = 0.010), the signal collapsed when the single gene HOXA10 was removed (p = 0.56) and disappeared when all detected HOX genes were included (p = 0.63); the HOXA cluster alone reversed direction (Supplementary Figure S1). We therefore do not treat a broad HOX positional program as a depot-identity axis; the defensible identity signal is carried by mesenchymal-patterning genes (TBX15/SHOX2) and by the VAT mesothelial–stromal program.

#### S-Results 2. Persistence of the depot signal after cell-composition and BMI/age adjustment

Because bulk depot differences can reflect cell-type proportions, we asked whether the SAT mesenchymal signal survived adjustment for mesothelial and macrophage signature scores. In a patient-random-intercept model, the depot effect on the mesenchymal module was largely retained after adjustment (~92% of the coefficient; $p < 0.0001$) (Figure S4). The depot effect also remained highly significant after adjustment for BMI and age (mesenchymal depot $\beta = -1.72$, $p < 0.0001$; mesothelial depot $\beta = +1.54$, $p < 0.0001$), indicating that depot-associated identity remained significant after adjustment for BMI and age in this cohort. We interpret the composition adjustment conservatively: the depot-associated mesenchymal signal is not fully explained by measured mesothelial and macrophage signatures. We do not claim independence from cell composition in general, as unmeasured fibroblast, vascular, and progenitor subpopulations may remain, and mesothelial and mesenchymal transcripts could in part co-originate from shared stromal populations [9,11].

#### S-Results 3. Depot-associated remodeling modules and adiposity (full detail)

The two depots were associated with distinct remodeling modules (Figure S6). SAT cross-sectional area correlated with the ECM cross-linking/stiffening module ($\rho = 0.77$, $p = 0.005$), whereas VAT area correlated with the vascular–hypoxia module ($\rho = 0.57$, $p = 0.013$) but not with macrophage content ($\rho = 0.03$, $p = 0.91$). However, because depot area was strongly collinear with BMI (SAT $\rho = 0.95$; VAT $\rho = 0.86$), we treated BMI not merely as a nuisance confounder but as an integrated marker of chronic energy surplus, and asked whether local area exerted an effect beyond overall adiposity. After BMI adjustment, the area–module associations did not persist (SAT ECM: partial $r = 0.08$, $p = 0.81$; VAT vascular–hypoxia: partial $r = 0.22$, $p = 0.38$). Because BMI and depot area were strongly collinear manifestations of adiposity, adjustment could not isolate an area-specific effect; the unadjusted associations are therefore interpreted as adiposity-linked depot remodeling rather than as effects uniquely attributable to local depot size, and the loss of area-specific significance reflects limited ability to separate global adiposity from local expansion in this cohort rather than absence of adiposity-linked transcriptional remodeling. The biologically important observation preserved by the data is that the two depots engage different remodeling modules as adiposity increases. Within SAT, LOX expression was higher in the obese than in the lean subgroup; because the SAT BMI distribution was bimodal (lean cluster 17–22 $kg/m^2$, obese cluster 30–36 $kg/m^2$, with no samples at 23–29—reflecting the surgical population; Section 2.1), we report this as a between-subgroup difference rather than a continuous dose–response relationship (Spearman $\rho = 0.93$ and Pearson $r = 0.83$ provided for transparency; Supplementary Figure S2).

#### S-Results 4. CD68 quantification in the histologic cohort

In an independent histologic cohort (n = 30; overlap with the RNA cohort = 1 patient), paired histology showed markedly higher interstitial fibrosis in SAT than in VAT (17.8% vs 4.2%, paired Wilcoxon $p \approx 3\times10^{-6}$) (Figure 3), providing independent, tissue-level support—rather than within-patient validation—for the SAT ECM/stiffening program. Representative sections illustrate more abundant collagen deposition in SAT relative to VAT (Figure 3A). The absolute CD68-positive area fraction remained very low in both depots. Although the median CD68-positive area fraction was modestly higher in VAT than SAT (VAT 0.031% [IQR 0.016–0.043] vs SAT 0.013% [IQR 0.009–0.031]; VAT > SAT in 22/30 patients; paired $p \approx 0.005$), the overall magnitude of CD68-positive staining was minimal in both adipose depots. Similar mean values were largely driven by a single SAT specimen with markedly increased CD68 positivity (Supplementary Table S3). Given the small absolute magnitude of CD68-positive staining and the absence of any association between VAT area and the macrophage RNA module, we did not interpret CD68 as strong histologic support for a VAT macrophage remodeling axis; the quantitative CD68 analysis (log scale) is shown in Supplementary Figure S5.

#### S-Results 5. Storage-versus-scaffold: robustness, storage-diversion, and CT attenuation (HU)

Because the histologic cohort had complete paired CT depot areas and attenuation for all 30 patients (no missing values), we examined the relationship between fibrosis density and depot size directly (Figure 5). Fibrosis fraction was inversely related to depot area in SAT ($\rho = -0.45$, $p = 0.012$) and, more weakly, in VAT ($\rho = -0.31$, $p = 0.09$): larger depots had lower fractional fibrosis. Strikingly, SAT specimens with high fibrosis had significantly smaller SAT area than low-fibrosis specimens (median 111 vs 162 $cm^2$, $p = 0.003$), consistent with a scaffold-associated ceiling on subcutaneous expansion. Despite the higher fractional fibrosis in SAT, the two depots differed sharply in scaffold economy (all measures are 2-D area-weighted CT–histology indices, not tissue-volume or collagen-mass measurements; Methods 2.6). The area-weighted fibrosis index (area × fraction) was higher in SAT than VAT (SAT > VAT in 27/30 patients; paired $p < 0.0001$), and the storage-to-scaffold index (area per percentage-point of fibrosis) was far higher in VAT (median 46.5 vs 8.7 $cm^2$ per %; VAT > SAT in 28/30 patients; paired $p < 0.0001$)—that is, omental VAT showed several-fold more adipose area per unit of fractional fibrosis than SAT (a between-depot ratio invariant to the percentage/fraction convention). Finally, higher SAT fibrosis showed a weak positive association with the VAT/SAT area ratio ($\rho = +0.36$, $p = 0.053$; BMI-adjusted partial $r = +0.33$, $p = 0.08$), in the direction expected if a scaffold-constrained SAT diverts storage toward the visceral compartment; however, this association was of borderline significance and did not persist after simultaneous adjustment for BMI, age, sex, and diabetes (partial $r = +0.19$, $p = 0.32$), and SAT fibrosis was not directly correlated with VAT area ($\rho = -0.05$). We therefore present the storage-diversion idea only as a hypothesis consistent with the direction of the data, not as an established relationship. In contrast, the inverse SAT fibrosis–area relationship was robust to outlier removal ($\rho = -0.44$ to $-0.49$ across trims) and held in the male and non-diabetic subgroups, and the storage-to-scaffold difference was robust to removal of the most extreme value (VAT > SAT in 27/29; $p < 10^{-6}$). Beyond fibrosis, CT attenuation (HU) behaved as a depot-associated imaging phenotype distinct from histologic fibrosis in the histologic cohort (complete for all 30 patients; Supplementary Figure S8). Attenuation was strongly and inversely related to depot area in both depots (SAT $\rho = -0.80$, VAT $\rho = -0.79$; both $p < 0.001$), and this relationship persisted after statistical adjustment for BMI in this cohort (SAT partial $r = -0.81$; VAT $r = -0.63$). SAT attenuation was lower (more negative) than VAT (median −93.7 vs −91.5 HU; SAT lower in 20/30; $p = 0.006$). In contrast, attenuation was only weakly associated with fibrosis fraction (SAT $\rho = 0.27$, $p = 0.16$; VAT $\rho = 0.33$, $p = 0.07$) and did not become a positive fibrosis surrogate after adjustment for depot size and BMI. We therefore interpret HU cautiously: it captures a depot-associated imaging phenotype distinct from histologic fibrosis, but its biological basis (e.g., adipocyte lipid fraction versus partial-volume, perfusion, or water content) and its sensitivity to ROI segmentation remain to be established, and all HU findings are reported as exploratory (Supplementary Table S3, Figure S8).

#### S-Results 6. Single-nucleus cell-of-origin and BMI associations (full detail)

At single-nucleus resolution, fibrillar collagen expression (COL1A1, COL3A1, COL6A3) was concentrated in adipose stem/progenitor cells (ASPCs), whereas LOX was distributed across stromal and vascular cell types (Figure S9). Within ASPCs, collagen and developmental-identity scores were higher in SAT than VAT (rank-biserial −0.71, $p = 0.010$ and −0.76, $p = 0.006$), and LOX showed a consistent SAT-high trend in paired ASPCs, although the paired comparison was underpowered (5/6 donors SAT > VAT; paired Wilcoxon $p = 0.063$). Mesothelial identity and monocyte-recruitment/macrophage signals were VAT-selective across cell types. Exploratory donor-level analyses provided convergent evidence that selected ECM and angiogenic programs increase with BMI, particularly in VAT (adipocyte ECM cross-linking versus BMI, VAT $\rho = 0.72$, nominal $p = 0.019$; VAT endothelial angiogenesis $\rho = 0.64$, nominal $p = 0.048$); these associations were based on ~9–10 donors and require validation in larger cohorts. Thus the public datasets validate the depot architecture, whereas our cohort uniquely connects that architecture to BMI, CT adiposity, attenuation, and histologic fibrosis.

#### S-Results 7. Future directions

Future directions. Several targeted additions would strengthen the model. Histologic confirmation of the VAT vascular–hypoxia program (e.g., CD31/PECAM1 endothelial density and HIF1A immunostaining) would give VAT the kind of tissue-level evidence that Masson trichrome provides for the SAT ECM program, which is presently asymmetric between depots. A fibrosis classifier trained on both depots, with blinded pathologist concordance scoring, would confirm the magnitude of the SAT–VAT fibrosis difference and the storage-to-scaffold indices. Reference-based deconvolution against a human adipose single-cell atlas would complement the marker-based cell-composition adjustment used here. Single-nucleus RNA sequencing and spatial transcriptomics would resolve whether depot-identity and remodeling signals arise from adipocytes or from stromal, vascular, and mesothelial compartments [12,13], directly addressing the composition caveat. DNA methylation or chromatin-accessibility profiling of paired, cell-fractionated specimens would test whether depot identity is epigenetically encoded, and in vitro perturbation (e.g.,

TBX15 or LOX manipulation in preadipocytes or explants) would test whether the identity–ECM coupling is causal. A longitudinal study relating baseline depot remodeling states to post-surgical weight loss and metabolic response is ongoing. Therapeutically, depot-specific ECM modulators (e.g., anti-fibrotic or LOX-targeted strategies) merit evaluation given the SAT stiffening program [17]. The single-nucleus deconvolution proposed here was performed against the Emont et al. atlas (Section 3.7), localizing fibrillar collagen expression to ASPCs; spatial transcriptomics and cell-fractionated profiling remain the natural next steps.

### Supplementary Tables

Table S1. Module gene definitions (as analyzed) and the mesothelial/immune/endothelial/erythroid exclusion list; per-variable data availability. (S1a demographics, S1b CT with HU quality flags, S1c module definitions.)

Table S1 (summary). RNA-seq cohort: 17 patients (sex M 16, F 1), 29 adipose specimens (SAT 11, VAT 18), including 10 within-patient SAT–VAT pairs. Age 65.0 [50.0–72.0] years; BMI 21.9 [20.5–31.5] $kg/m^2$ (bimodal: lean ~17–22 and obese ~30–40, few overweight). CT area/attenuation available for all; histology (Masson/CD68) was performed in a largely non-overlapping independent cohort ($n = 30$). Complete patient-level metadata are provided in Supplementary Table S1 (Excel).

Table S2. Paired SAT–VAT differential expression.

Table S3. CT–RNA associations (raw and BMI-adjusted), mixed-model coefficients (β, 95% CI, SE, p), and CD68 summary statistics (median [IQR], paired difference).

Table S4. Sample-flow accounting for depot assignment and library submission batches.

Table S5. Storage-versus-scaffold analysis (fibrosis–area relationships, area-weighted fibrosis index, storage-to-scaffold index with units, robustness, and HU associations).

Table S6. External validation: genome-wide depot log2 fold-changes and FDR in our cohort, GTEx, and the Emont single-nucleus atlas, with the 341 GTEx-replicated, directionally concordant DEGs and column definitions (SuppTable_ThreeLayer_validation.xlsx).

### Supplementary Figures

Figure S1. HOX analyses and non-robustness (single-gene dependence; full-HOX null).

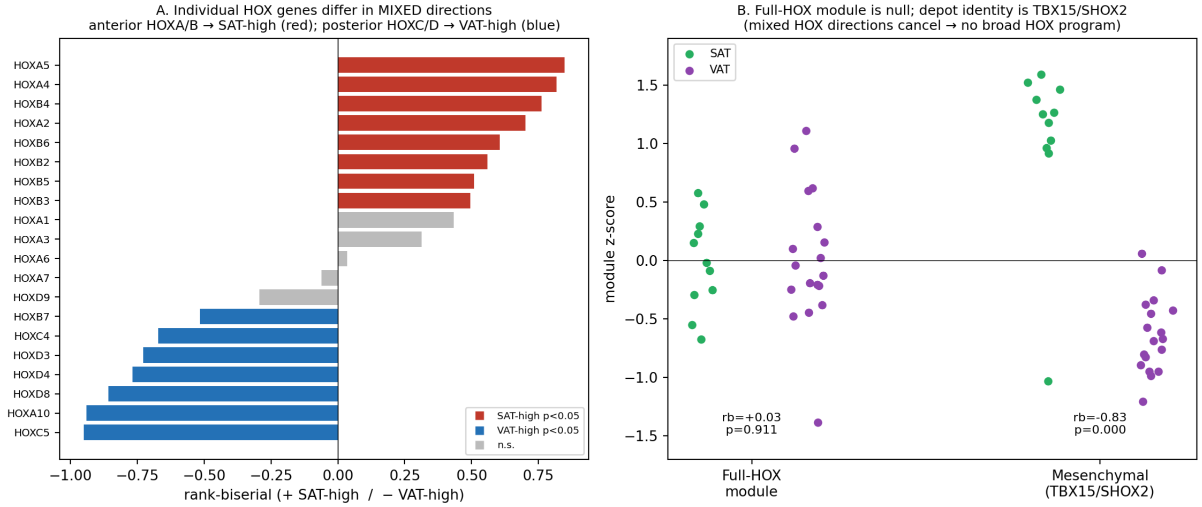


Figure S2. LOX–BMI with bimodal BMI distribution; Spearman/Pearson and subgroup comparison.

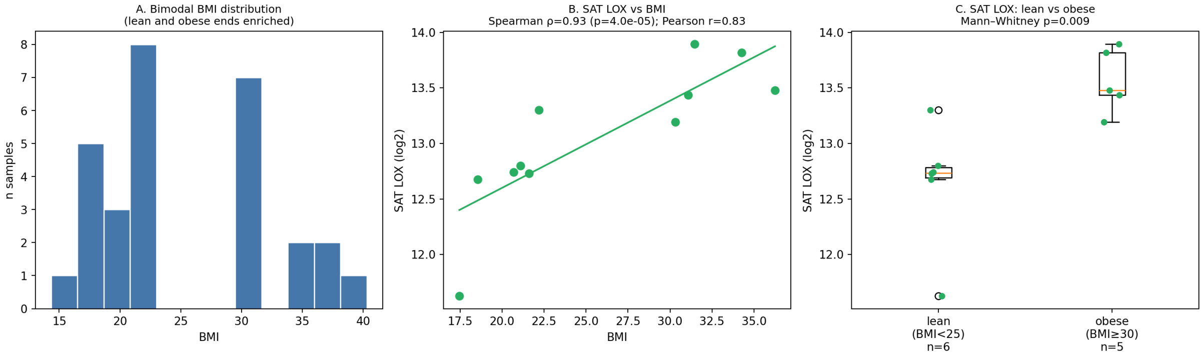


Figure S3. Leave-one-patient-out classifier with random-gene null distribution.

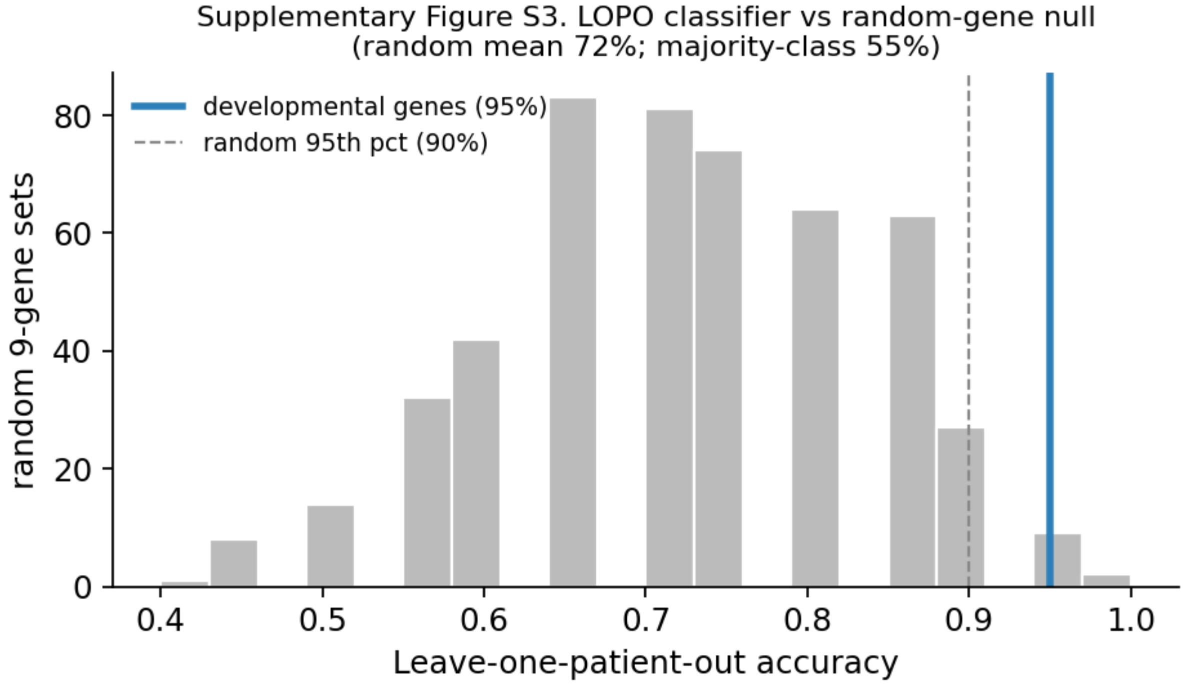


Figure S4. Persistence after adjustment for measured cellular signatures. Forest plot of the depot coefficient (VAT − SAT, standardized) for the mesenchymal module, unadjusted versus adjusted for mesothelial and macrophage scores in a patient-random-intercept model; points are coefficients, whiskers are 95% CI. Retained effect = 100 × |β_adjusted|/|β_unadjusted| ≈ 92% (n = 10 pairs; p < 0.0001 in both models).

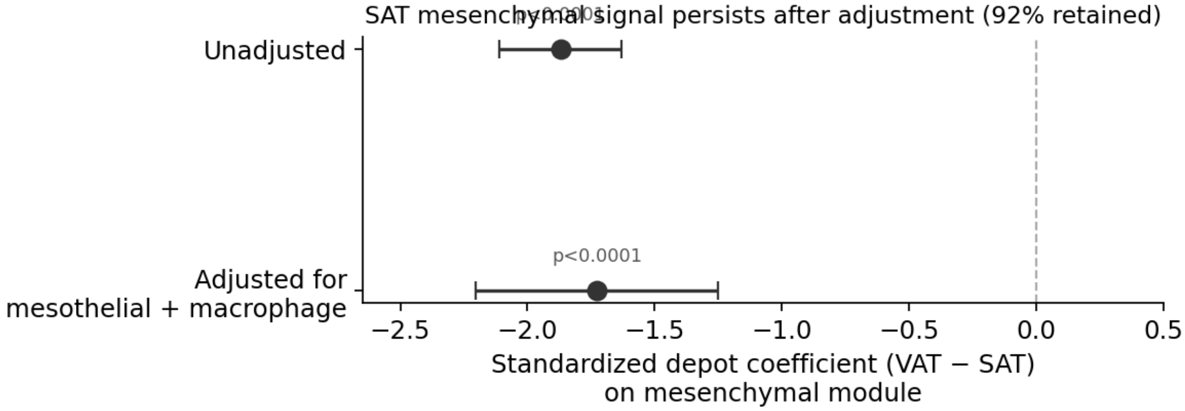


Figure S5. CD68 quantification (log-scaled) in the independent cohort. CD68-positive area fraction in paired SAT and VAT specimens (n = 30). Values are shown on a logarithmic scale because the absolute CD68-positive area fraction was very low in both depots. Although VAT showed a modestly higher median CD68-positive area fraction than SAT, CD68-positive staining remained sparse overall in both depots.

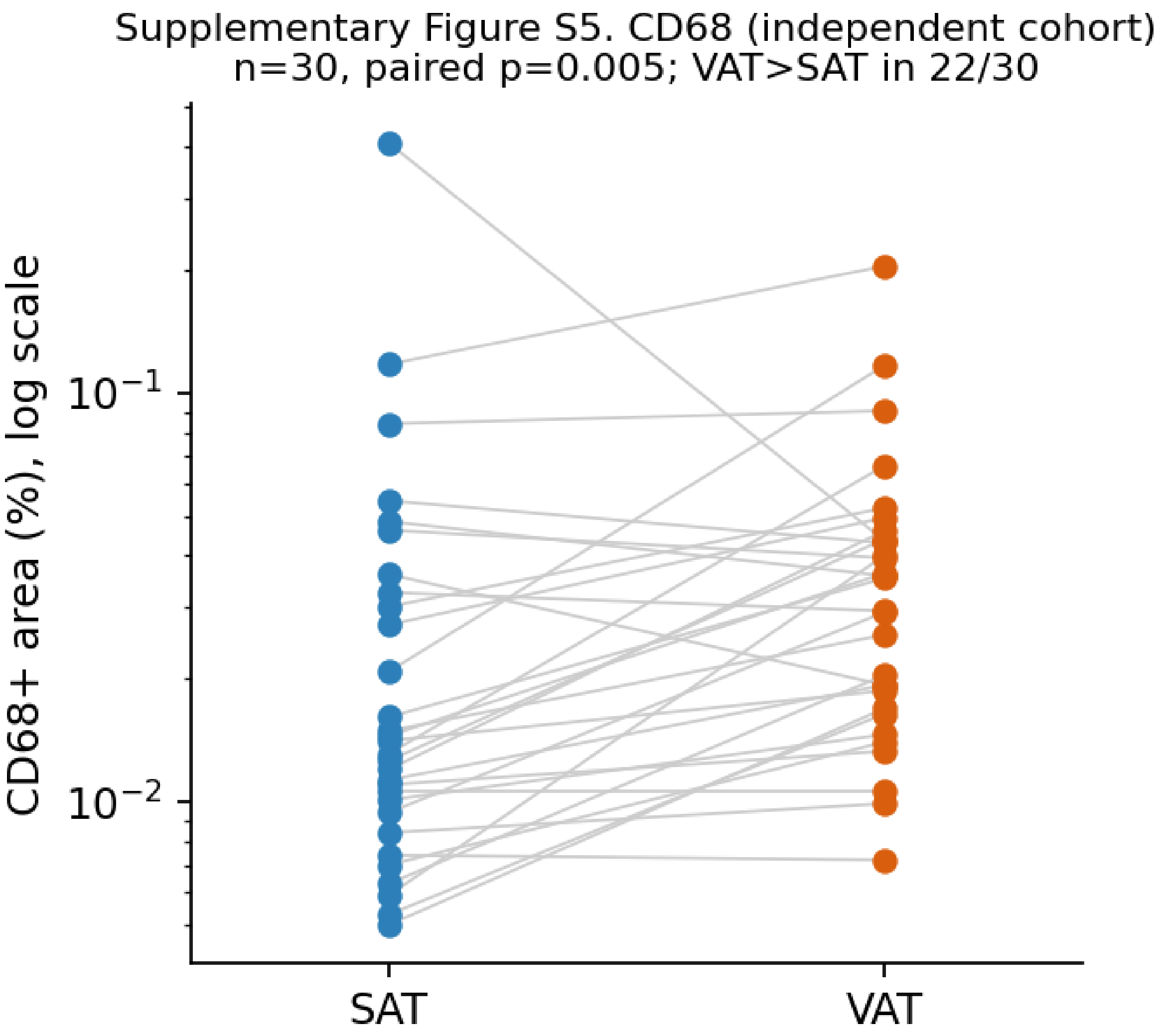


Figure S6. CT–RNA associations (baseline, RNA cohort). SAT area ↔ SAT ECM/stiffening module ($\rho$ = 0.77, p = 0.005, n = 11); VAT area ↔ VAT vascular–hypoxia module ($\rho$ = 0.57, p = 0.013, n = 18). Regression lines are for visualization only. These associations did not survive BMI adjustment (Section 3.4); depot area and BMI are strongly collinear.

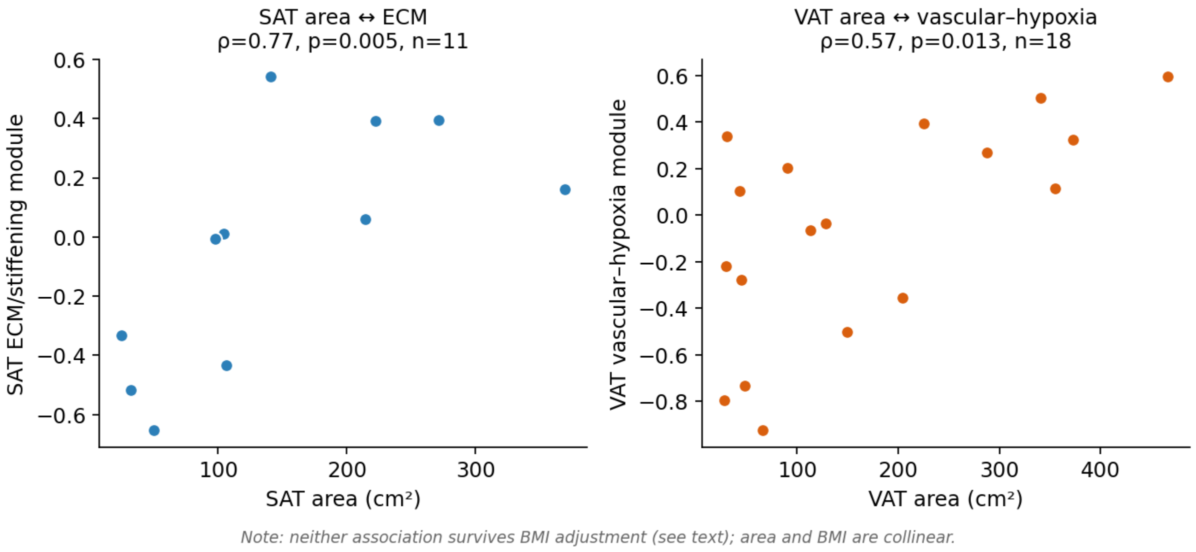


Figure S7. PCA sensitivity excluding the VAT outlier (separation retained).

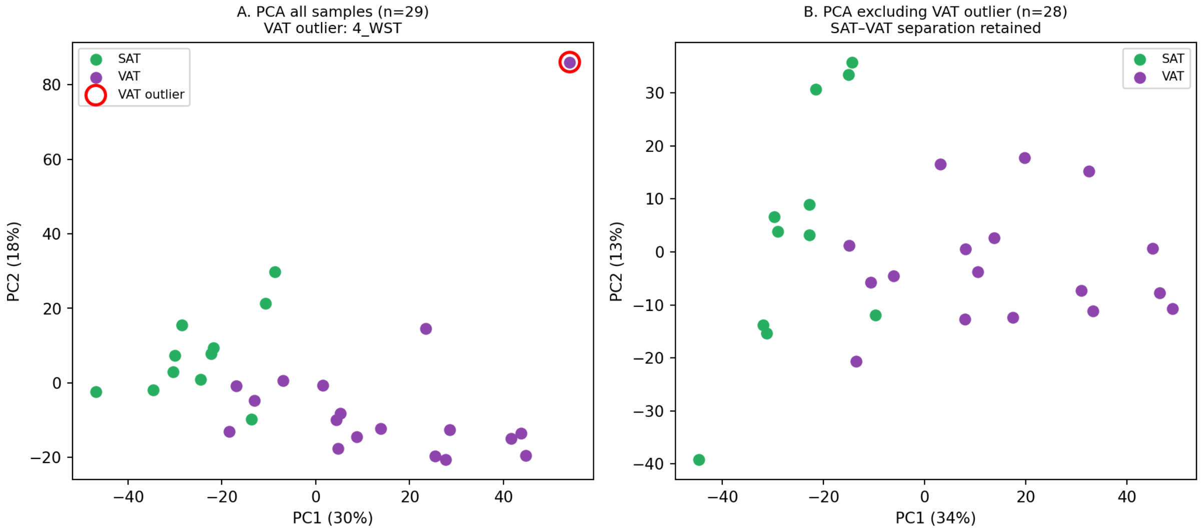


Figure S8. CT attenuation (HU) as a depot-associated imaging phenotype distinct from fibrosis: HU–area relationship, SAT vs VAT HU difference, and weak HU–fibrosis association; biological basis and segmentation sensitivity remain to be validated.

Supplementary Figure S7. CT attenuation is a depot-associated imaging phenotype distinct from fibrosis (biological basis and segmentation sensitivity require validation) (n=30)

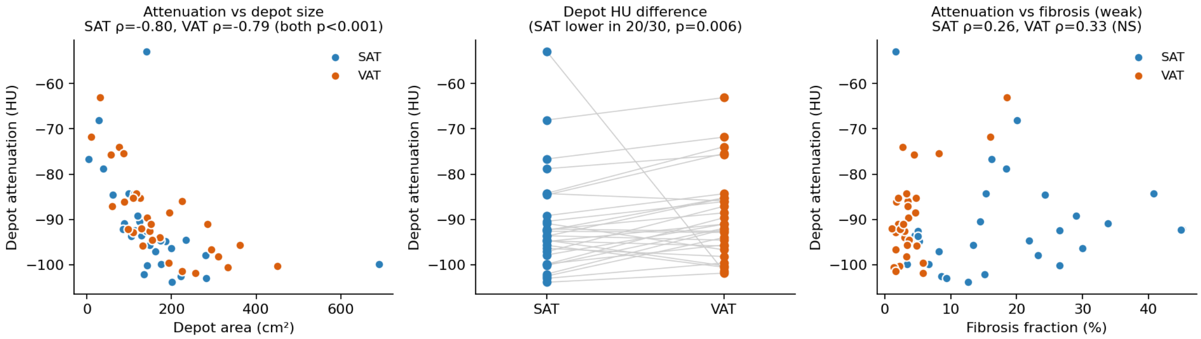


Figure S9. Single-nucleus context for depot ECM programs (Emont et al. atlas [13]; 137,684 nuclei). (A) Cell-type expression of ECM/stiffening genes: fibrillar collagens (COL1A1/COL3A1/COL6A3) are concentrated in ASPCs. (B) LOX expression by cell type and depot, distributed across stromal and vascular cells rather than ASPC-exclusive. (C) Within-cell-type depot contrasts (rank-biserial). (D) Adipocyte ECM cross-linking versus BMI (exploratory; nominal P values; ~9–10 donors). (E) Depot module means on true tissue-level pseudobulk (all nuclei summed per donor × depot). (F) Cross-dataset concordance summary.

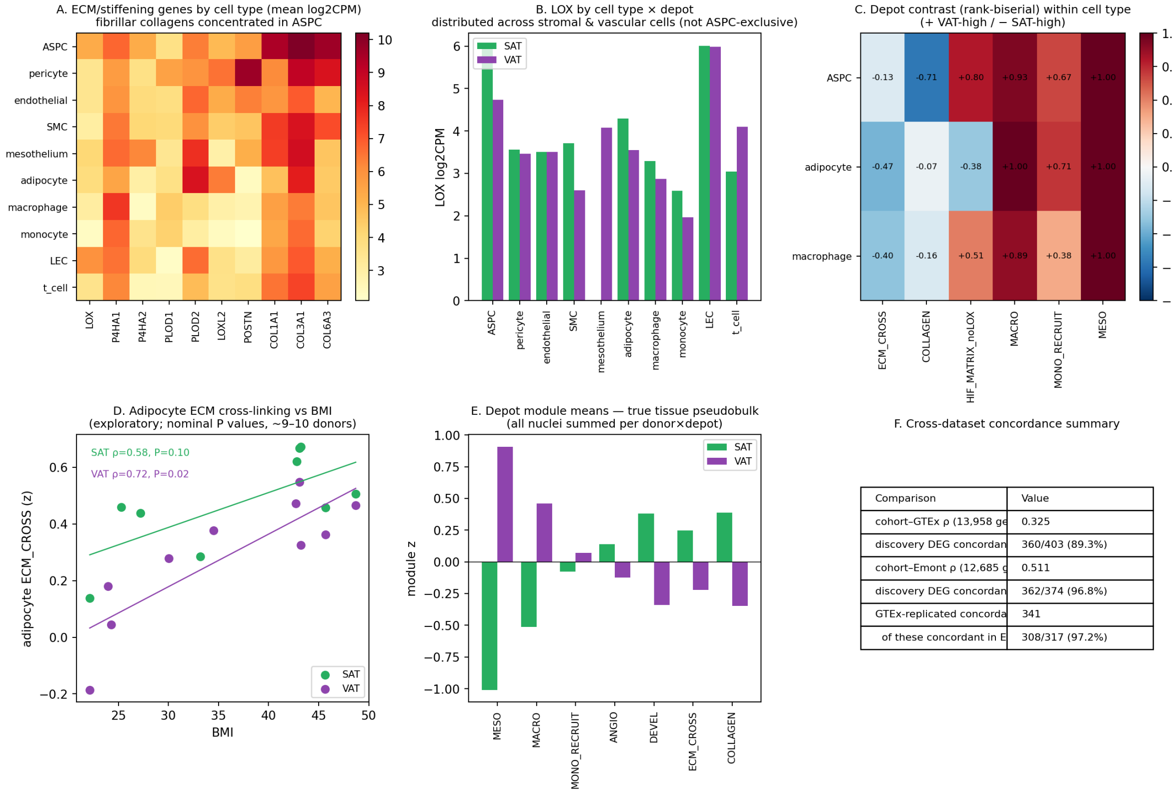


| Comparison | Value |
|---|---|
| cohort–GTEx ρ (13,958 ge | 0.325 |
| discovery DEG concordan | 360/403 (89.3%) |
| cohort–Emont ρ (12,685 g | 0.511 |
| discovery DEG concordan | 362/374 (96.8%) |
| GTEx-replicated concorda | 341 |
| of these concordant in E | 308/317 (97.2%) |
